# Structure of the human HIRA histone chaperone with a nucleosome suggests a stepwise nucleosome assembly mechanism

**DOI:** 10.64898/2026.05.17.725740

**Authors:** Wei Tian, Siyu Chen, Liqi Yao, Vignesh Kasinath, Karolin Luger

## Abstract

The restoration of chromatin in the wake of a DNA or RNA polymerase is essential to maintain the integrity of eukaryotic genomes. Human HIRA is a 1.8-megadalton, three-subunit histone chaperone that mediates all replication-independent deposition of the histone variant H3.3 at active chromatin regions^1–6^. Disruption of HIRA perturbs active-chromatin organization and has wide-ranging consequences for development, cellular senescence, and genome integrity^7–11^. Despite its central biological role in reassembling nucleosomes post-transcription, the structure of native human HIRA and the mechanism by which it organizes histones and DNA during nucleosome assembly remain unknown. In particular, the function of the largest HIRA subunit CABIN1 is enigmatic. Here, we show that HIRA is not simply a passive histone hand-off factor but remains engaged across multiple stages of nucleosome assembly, including a close interaction with the nucleosome. Cryo-EM structures reveal that HIRA forms an extended arch-like structure that binds the nucleosome primarily through extensive CABIN1 contacts with histones, histone tails, nucleosomal DNA, and linker DNA, during the final stage of nucleosome assembly. Together, our results suggest a testable mechanism for HIRA-mediated nucleosome assembly and product release and provide the basis for elucidating the molecular details of this fundamental biological process.

## Main

Nucleosome assembly is essential for restoring and maintaining chromatin organization during DNA replication, transcription, and repair^12^. This process occurs through a stepwise pathway in which (H3-H4)_2_ tetramer is first deposited onto DNA, followed by sequential incorporation of two H2A-H2B dimers to generate a hexasome intermediate and ultimately a complete nucleosome where 147 base pairs of DNA are tightly wrapped around the H3-H4-H2A-H2B octameric core. This pathway is driven by histone chaperones that ensure ordered and well-timed deposition of histones and histone variants^13,14^. Histone chaperones are highly diverse and generally share little sequence or structural similarity. Different chaperones are specialized for in distinct histone classes, histone variants, or chromatin assembly pathways, allowing cells to route specific histones to defined genomic regions and biological contexts. Along these pathways, histones are often passed from one chaperone to another in a coordinated “shuffle,” frequently linked to histone post-translational modifications that help specify their assembly. Histone chaperones have long been viewed as passive escorts of histones through individual steps of nucleosome assembly with transient product engagement at most^14^.

Human HIRA is the major replication-independent histone chaperone complex responsible for histone variant H3.3 deposition during active transcription. It is composed of three large subunits that assemble into a massive complex (see Fig. 2b): HIRA_s_ (111.8 kDa; the subunit for which the entire complex is named), UBN1 (121.5 kDa); and CABIN1 (246.4 kDa). Although the precise stoichiometry of the human complex has not been fully established, it is thought to contain six HIRA_s_ subunits, two CABIN1 subunits, and four UBN1 subunits^15,16^. The small H3.3-H4 chaperone Asf1a initiates HIRA-dependent nucleosome assembly by specifically delivering H3.3 to the HIRA complex through interactions with HIRA_s_ and UBN1 subunits^17,18^. Previous studies have assigned different roles to the three HIRA subunits. Six HIRA_s_ subunits form the core of the complex, by engaging UBN1 with its N-terminal WD40 region^15,19^. A small region adjacent to the WD40 domain binds Asf1a, and its C-terminal region assembles into two homo-trimers that bind CABIN1^16,17^. Trimerization and WD40 domains are important for full nucleosome assembly activity in cells^16,20^. UBN1 likely confers H3.3 specificity and DNA binding capability through its conserved Hpc2-related domain (HRD) and middle domain, respectively^18,21^. The function of CABIN1 is largely unknown beyond its potential role as a transcriptional corepressor^22,23^.

There is a major gap in our understanding of how the native human HIRA complex binds and organizes histones and DNA to mediate nucleosome assembly. Most published biochemical studies had to rely on isolated protein subunits or domains, leaving the architecture and mechanism of the full complex unresolved. A recent structure of the yeast homolog Hir, bound to Asf1 and histones H3-H4, provided a first view of the overall stoichiometry and architecture during an early assembly step^15^. However, yeast Hir and human HIRA differ substantially in subunit composition and size, and many questions remain with respect to the entire nucleosome assembly reaction. Unlike metazoans, yeast lacks the histone variant H3.3 and as such the biological function of yeast HIRA might differ from metazoan HIRA^24^.

Nucleosome assembly is an essential process in all eukaryotes, yet mechanistic and structural insight into how this complex, multi-step task is orchestrated in an ATP-independent manner is sparse. Our biochemical and structural investigations of the massive HIRA complex in the final stage of nucleosome assembly provide first insights into the mechanism of this important nucleosome assembly factor that maintains chromatin structure during active transcription.

## Results

### HIRA is involved in all steps of H3.3-containing nucleosome assembly

We were able to purify the human HIRA complex (subsequently referred to as HIRA) to homogeneity upon overexpression in a human embryonic kidney (HEK) 293 GnTI^-^ cell line (Fig. 1a). Since HIRA is responsible for histone variant H3.3 deposition, we first compared HIRA binding affinity for H3.3-H4 and H3.1-H4 via fluorescence polarization (FP). Although a small fragment of UBN1 in isolation was previously shown to distinguish between the two histone complexes, our data reveals that the HIRA complex binds both H3.3 and H3.1 histone complexes with an equal affinity of ≤ 0.5 nM (Fig.1b)^18^. HIRA also binds histones H2A-H2B, albeit with slightly lower affinity (∼3 nM; Fig. 1b), and interacts with DNA (Fig. 1e, lane 3). These results are consistent with published data showing that the isolated UBN1 middle domain binds DNA, and that yeast CABIN1 contains extensive basic regions^15,21^. Surprisingly, HIRA binds to fully assembled nucleosomes with high affinity, even in the absence of linker DNA (Fig. 1c). This is unexpected as most histone chaperones facilitate nucleosome assembly by masking the DNA-binding surface on histones, and do not interact strongly with the final nucleosome product^25,26^.

**Figure 1:**
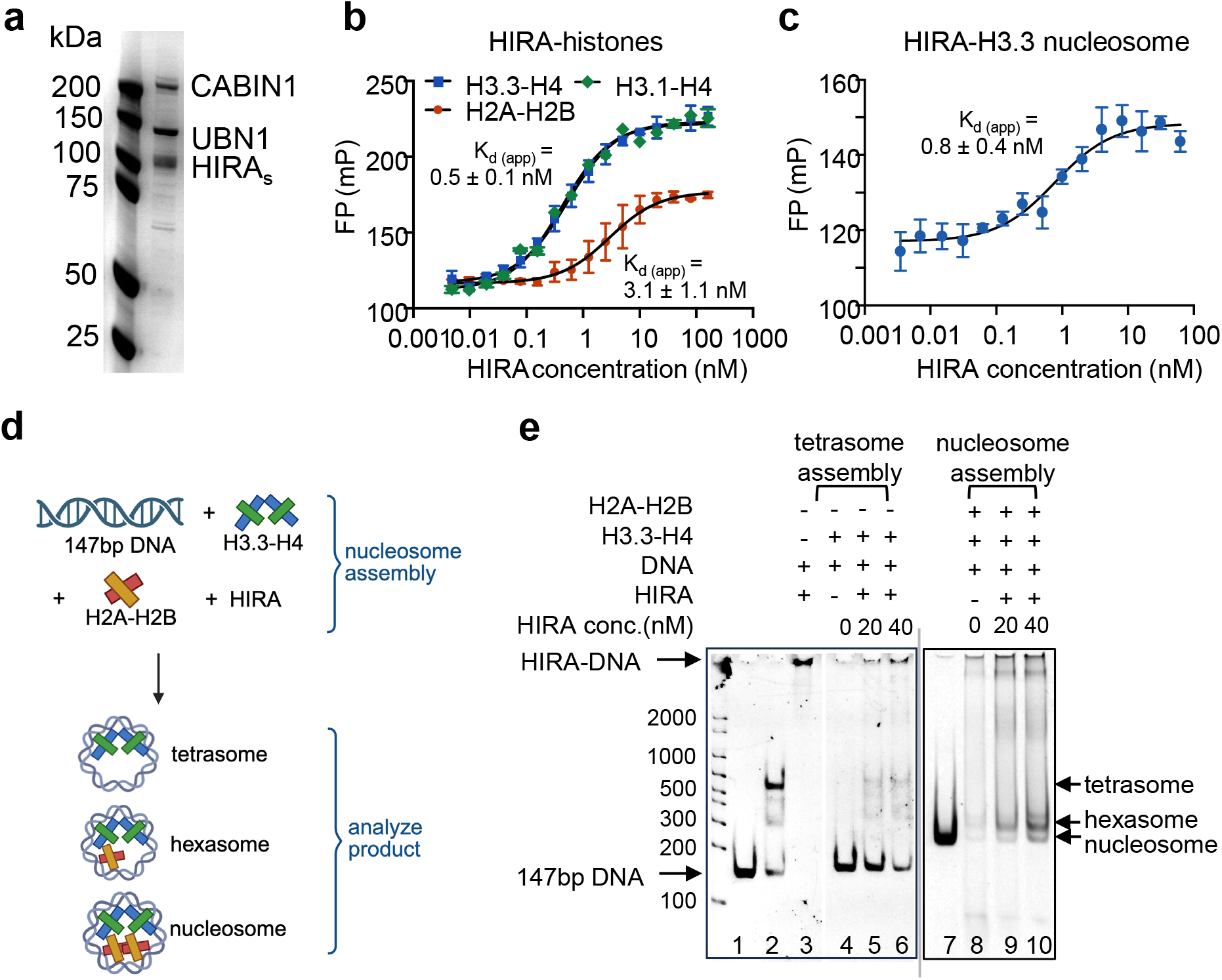
HIRA is involved in the entire nucleosome assembly process. **(a)** HIRA complex purified from mammalian cells (SDS-PAGE). **(b)** Fluorescence polarization assay of HIRA with fluorescently labeled histones. Kd and standard errors (s.e.) were determined from three replicates. **(c)** Fluorescence polarization assay of HIRA with Alexa-488 labeled nucleosomes (assembled with 147 bp DNA). Kd and standard errors (s.e.) were determined from three replicates. **(d)** Schematic of HIRA-dependent nucleosome assembly assay. **(e)** HIRA is active in nucleosome assembly. 5% native gel (stained with ethidium bromide (EtBr)). Lane 1: free 147 bp DNA, lanes 2 and 7: pre-assembled tetrasome and nucleosome controls, with a presumed ditetrasome band appearing in the reconstituted tetrasome sample. Lane 3: HIRA binds 147 bp DNA, trapping it in the well. Lanes 4-6: H3.3-H4 mixed with DNA with the absence and presence of increasing amounts of HIRA, as indicated. Lanes 8-10: H2A-H2B dimer was added to reactions from lanes 4-6. Tetrasome, hexasome and nucleosome products are formed as indicated.

The nucleosome assembly activity of HIRA has been qualitatively demonstrated using *Xenopus laevis* egg extracts with plasmid DNA as the substrate. However, the complexity of extract-based systems and the use of incompletely defined HIRA complex have limited the mechanistic characterization of HIRA-mediated nucleosome assembly^4,20^. Here we employ an established nucleosome assembly assay with purified components^27^ (workflow shown in Fig. 1d). We observe the HIRA-dependent formation of a tetrasome product (a nucleosome lacking both H2A-H2B dimers; Fig. 1e, lanes 5-6). Upon addition of H2A-H2B dimers, hexasomes (nucleosomes lacking one H2A-H2B dimer) as well as complete nucleosomes are formed in a HIRA-dependent manner (Fig. 1e, lanes 9-10). We further tested whether HIRA promotes H2A-H2B deposition by adding pre-formed HIRA-H2A-H2B to tetrasomes resulting in a HIRA-dependent increase in hexasome and nucleosome (Supplementary Fig. 1b). Our results suggest that purified recombinant HIRA functions in all steps of nucleosome assembly.

### Overall architecture of the HIRA-nucleosome complex

The interaction between HIRA and the nucleosome might provide insight into the mechanism of assembly. To visualize the interaction, we prepared nucleosome-HIRA complexes for analysis by cryo-EM. These were assembled with nucleosomes reconstituted on either 147 bp or 207 bp ‘601’ positioning sequence DNA (the latter with 30 bp of linker DNA on both ends). To stabilize the complexes (subsequently referred to as HIRA-Mn147 and HIRA-Mn207, respectively), we performed GraFix followed by negative-stain EM, which indicated the presence of uniform particles ∼40 nm in size (Supplementary Fig. 2). Samples were then applied to EM grids with thin-carbon film (1-2 nm) and analyzed by cryo-EM. Initial 2D classification confirmed an elongated complex of ∼400 Å in length and revealed two nucleosomes bound on either end of the HIRA core (Fig. 2a).

**Figure 2:**
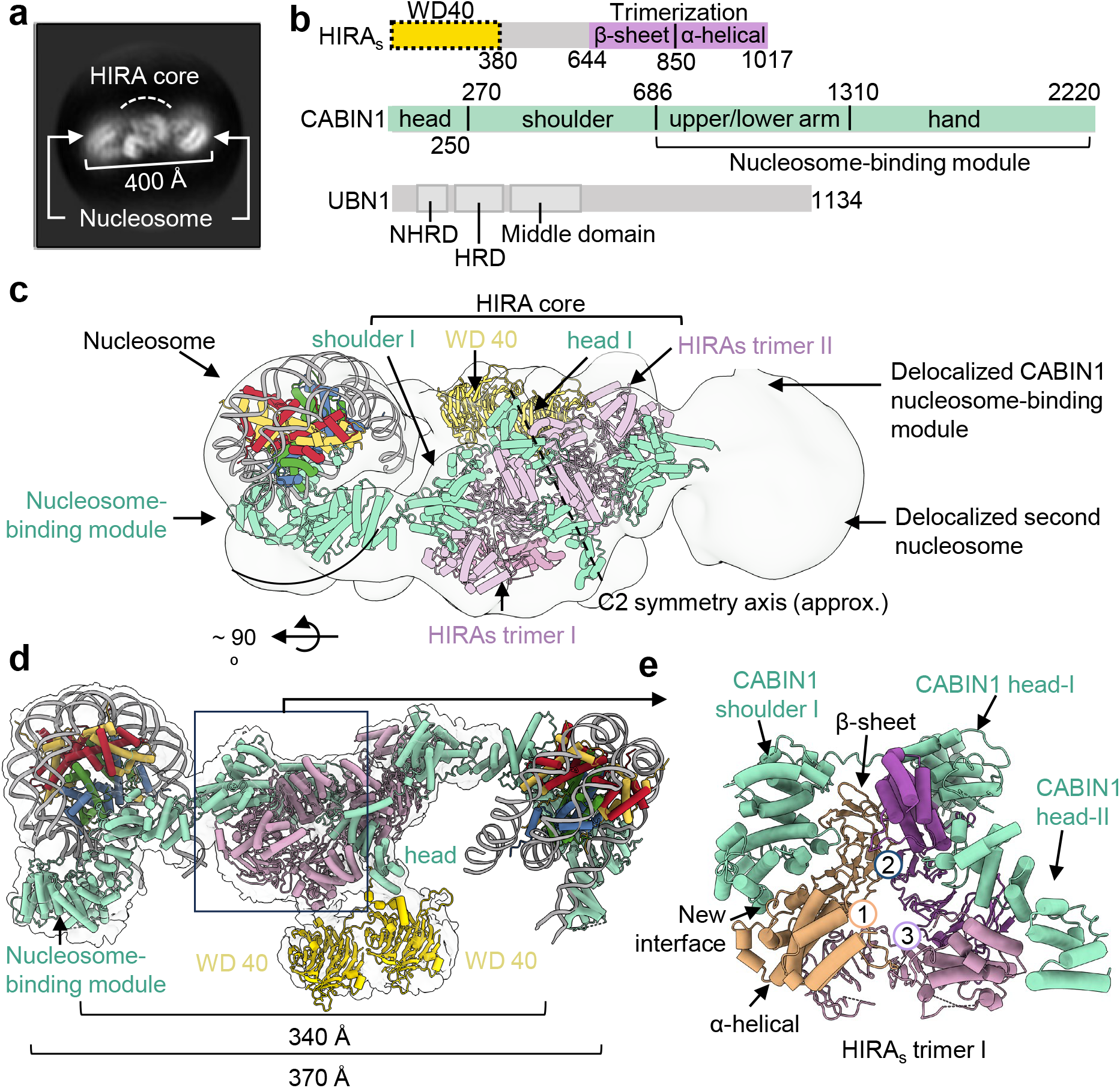
Overall architecture of HIRA-nucleosome complex. **(a)** Representative 2D class average from a cryo-EM data collection of the HIRA-nucleosome complex, showing two nucleosomes bound at opposite ends of the HIRA core. (**b)** Domain organization of HIRA subunits; only the colored regions were resolved in the cryo-EM map. **(c)** Low-resolution cryo-EM map showing the overall architecture of the HIRA-nucleosome complex. The experimentally determined models of the HIRA core and nucleosome-binding module of CABIN1, together with the nucleosome model, were rigid-body fitted into the map to visualize the complete complex. Density corresponding to the second nucleosome and the distal CABIN1 nucleosome-binding module is delocalized. **(d)** Overview of the entire HIRA-nucleosome complex based on C2 symmetry. The extrapolated distance between the two hand domains of CABIN1 is 340 Å, between two nucleosomes is 370 Å. **(e)** Enlarged view of the nucleosome-proximal HIRA trimer (Trimer I). The three HIRA trimerization-domain protomers are labeled 1, 2, and 3. The CABIN1 head occupies the dimerization interface formed by trimerization domains 2 and 3, whereas the CABIN1 shoulder associates with the α-helical subdomain of HIRA subunit 1 through an additional interface not observed in yeast Hir. The HIRA trimerization domain consists of a central β-sheet base and C-terminal α-helical subdomains.

Upon 3D analysis, density for one of the two nucleosomes was more clearly identified at the periphery of the structure and was used for focused refinement (see Methods for details). In the HIRA-Mn147 complex, additional density near the nucleosomal DNA was less defined, indicating that stable HIRA-nucleosome interaction benefits from the presence of linker DNA. Therefore, further analysis was focused on the HIRA-Mn207 complex.

The overall organization of human HIRA resembles the arch-shaped, C2-symmetric architecture reported for yeast Hir^15^, but with a much wider span (see below). The widest distance spanned by the HIRA complex is ∼340 Å, and the distance between two nucleosomes held at the endpoints of the arch is ∼370 Å (Fig. 2a, c, and d). The apex of the arch is formed by the two well-defined homo-trimeric assemblies of the HIRA trimerization domain (Fig. 2b-e, Supplementary video 1). These are connected by two copies of CABIN1, the largest HIRA subunit, which is predicted to adopt an extended α-solenoid architecture enriched in TPR-like repeats, similar to what has been described for its yeast homolog Hir3^15,23^. Two CABIN1 subunits make up the arms of the arch and each clutches one nucleosome at its end (Fig. 2d). For ease of description, we subdivided CABIN1 into ‘head, shoulder, upper arm, lower arm, and hand’ regions (Fig. 2b). In this terminology, the head and shoulders of CABIN1 hold the two homotrimers of HIRAs together, while the upper arm interacts with the histone surface. The lower arm clutches the nucleosomal superhelix, and the distal hand is positioned to guide the exiting DNA (Fig. 2d, and see below). The upper and lower arm and hand comprise the nucleosome binding module. We also observed density connected to the CABIN1 N-terminal domain and the HIRA trimerization domain on one side of the complex, which we interpret as two HIRAs WD40 domains (Fig. 2d). The UBN1 subunits and the remaining four WD40 domains were not observed in our structure.

The N-terminal head domains of the two CABIN1 molecules connect the two trimers of HIRAs, and one monomer in each trimer is additionally bound by a ‘shoulder domain’ (Fig. 2e). The HIRA trimerization domain itself can be divided into a β-sheet subdomain and a C-terminal α-helical subdomain (Fig. 2b, 2e). β-sheet elements from the three subunits form the central trimeric base, while the α-helical subdomains are positioned peripherally and provide additional interacting surfaces for the CABIN1 shoulder.

The two trimers in the HIRA core are structurally nearly identical (Supplementary Fig. 3a). In yeast, an analogous central core is assembled from two Hir1-Hir2-Hir2 heterotrimers, held together by the Hir3 (yeast CABIN1 equivalent) head region^15^. We compared the HIRA trimer architecture in our complex with the corresponding region in yeast Hir and with the crystal structure of the human HIRA_s_ trimer (Supplementary Fig. 3b-c). Across the three structures, the central HIRA_s_ β-sheet bases superpose closely, indicating that this framework is structurally rigid. The orientation of the α-helical subdomains is more variable, suggesting context-specific displacement.

### CABIN1 engages the nucleosome through multivalent interactions with histones and DNA

The observed extensive interactions between the CABIN1 arm and the nucleosome (Fig. 3) are unexpected and unusual since histone chaperones usually do not bind their assembly product. The CABIN1 upper arm approaches the histone octamer near the acidic patch (formed by histones H2A and H2B) and the H4 N-terminal tail, and makes extensive interactions with the nucleosomal DNA. To validate interaction interfaces, we performed crosslinking-mass spectrometry (XL-MS; Supplementary Fig. 4a, b). A CABIN1 region (amino acids 380-420) from within the shoulder region crosslinks to the acidic patch, as well as to the N-terminal tail of histone H4 (Supplementary Fig. 4). However, alphaFold predictions suggest that a short α-helix encompassing this region (amino acids 408-419) and adjacent loops extend away from the shoulder domain into the nucleosome-facing surface of the upper arm domain (Supplementary Fig. 4c). We were able to build this portion of CABIN1 within the nucleosome-focused map (3.4 Å) as well as within a lower-resolution reconstruction of CABIN1 alone (∼6.9 Å) with reasonable confidence. The side chains of R392 and R396 insert into the nucleosomal acidic patch, with R396 assuming the role of the ‘arginine anchor’ observed in all acidic-patch interactors^28^ (Fig. 3a-b). The nucleosome-interacting CABIN1 feature extending from the shoulder to integrate into the upper arm appears to be conserved in yeast (Supplementary Fig. 5).

**Figure 3:**
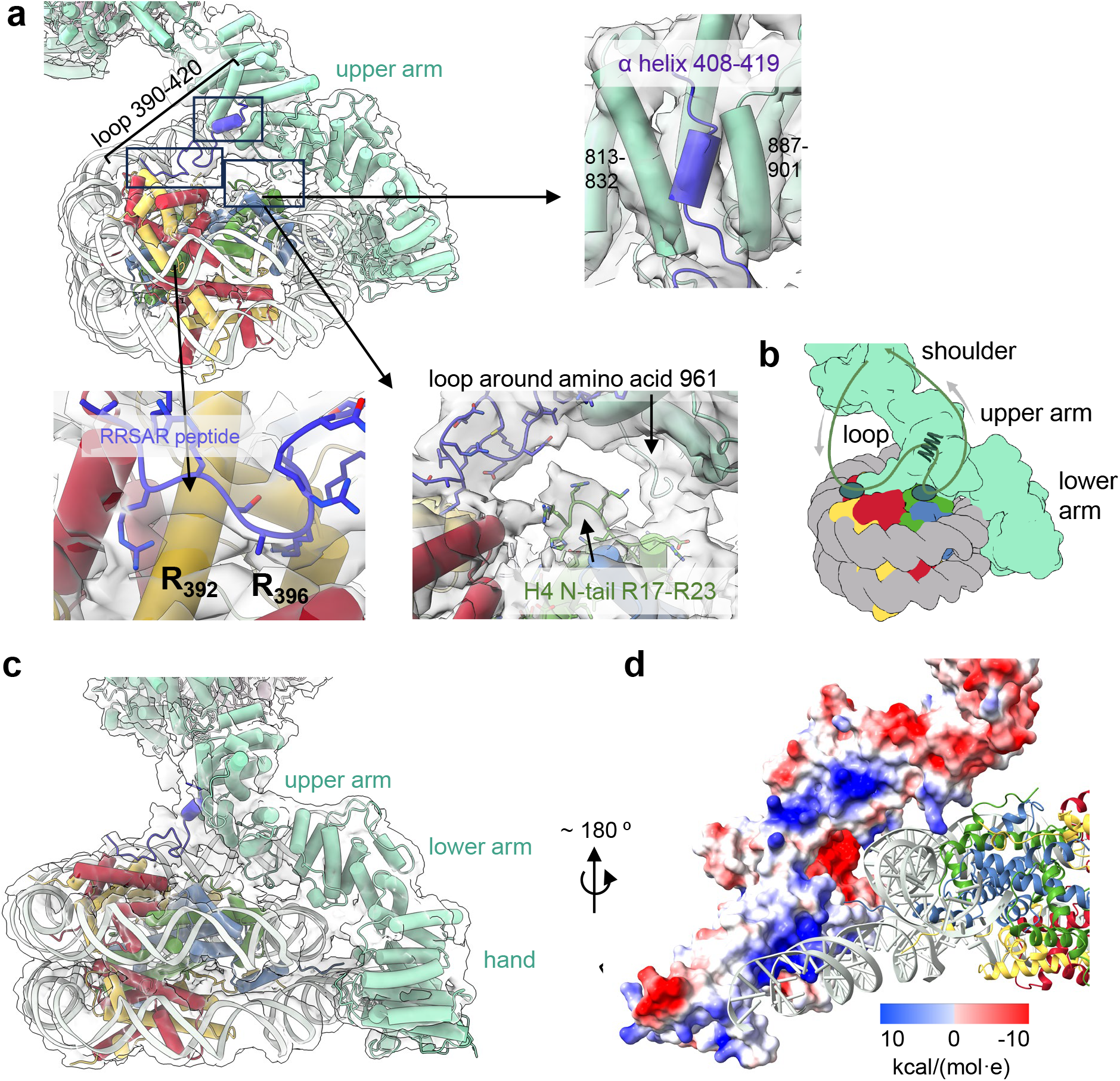
CABIN1 engages the nucleosome through multivalent interactions. **(a)** Overview of CABIN1 engaging the nucleosome. Nucleosomal DNA is shown in grey, H2A in yellow, H2B in red, H3.3 in blue, H4 in green and CABIN1 in light green. The CABIN1 nucleosome-interacting regions are highlighted in light blue. Zoomed-in regions of interactions between histones and CABIN1 are shown in three insets. **(b)** Simplified representation of CABIN1-nucleosome interactions, same orientation as in (a). **(c)** CABIN1 lower arm and hand domains clasp both gyres of the nucleosomal superhelix (spanning SHL +1 and -7). **(d)** Electrostatic surface of the CABIN1 lower arm and distal hand region, guiding the linker DNA. Inset shows Coulombic electrostatic potential in kcal/(mol·e).

In most nucleosome structures, the H4 N-terminal tail is flexible and can only be observed from R23 onwards. In our map, we can assign density corresponding to H4 residues R17-R23, directed toward the CABIN1 390-420 loop. This interaction is corroborated by XL-MS (Fig. 3a, b and Supplementary Fig. 4a, b). Additionally, a loop around amino acid 961 is poised to stabilize the N-terminal tip of the H4 α1 helix. Intriguingly, neither XL-MS nor our model indicates any interactions between the entire HIRA complex and H3.3 in this context.

In addition to the contacts with the acidic patch and the H4 tail, human CABIN1 also extensively engages both gyres of the nucleosomal DNA as well as linker DNA through its lower arm and hand region (Fig. 3c, d). Positively charged side chains in the CABIN1 lower arm are positioned near the central region (SHL1) of the nucleosomal DNA (Supplementary Fig. 6). The distal region of the hand domain is positioned the final turn of nucleosomal DNA (SHL-7), effectively stapling the two DNA gyres together (Supplementary Fig. 6). Additional TPR-like repeats in the hand domain then guide the linker DNA in a straight trajectory via a continuous basic surface (Fig. 3d). Notably, this positively charged track is also present in yeast Hir3 and *Chaetomium thermophilum* Hir3^15^.

Electrostatic surface analysis of the full CABIN1 model reveals two additional positively charged patches that in our model do not engage DNA (Supplementary Fig. 7a-c). One is located on the backside of the hand domain and is also observed in the yeast Hir3 structure^15^. In yeast Hir3, it forms a positively charged channel with a central opening speculated to bind single-stranded DNA or RNA^15^, whereas the human CABIN1 structure displays a continuous basic surface (Supplementary Fig. 7d). The second patch is positioned adjacent to the DNA binding region contacting SHL1 (Supplementary Fig. 7b). We also note a pronounced negatively charged band along the backside of the upper and lower arm (Supplementary Fig. 7e). Although the function of the intriguing charge distribution (apart from the established DNA binding regions) is unknown, their arrangement invites speculation on the mechanism by which HIRA assembles nucleosomes (see discussion).

### CABIN1 arm plasticity enables HIRA to accommodate different chromatin substrates

To address to what extent the CABIN1 arms contribute to the ability of the HIRA complex to bind products and substrates, we performed two sets of structural comparisons: first, a cross-species comparison between human HIRA-Mn207 and yeast Hir-Asf1-H3-H4 (PDB 8GHN)^15^, and second, a substrate-state comparison between human HIRA-Mn207 and HIRA in complex with a di-nucleosome construct, where the two nucleosomes are connected by 60 bp DNA (subsequently referred to as HIRA-Di60).

We superimposed the HIRA-Mn207 complex onto the yeast Hir-Asf1-H3-H4 complex using HIRA_s_ trimer I as the reference (Fig. 4a). The CABIN1 arms of human HIRA adopt a markedly more open conformation than in the yeast complex, with the distance between the two CABIN1 hands measuring 340 Å versus 130 Å in yeast. To assess whether this is caused by sequence differences between human HIRA and yeast Hir, or rather induced by the different binding partners (nucleosome versus Asf1-H3-H4), we prepared negative-stain grids of the human HIRA-Asf1a-H3.3-H4 complex (Supplementary Fig. 8a-b). The overall dimensions of this complex resemble those of the HIRA-Mn207 complex (340-370 Å), indicating that the open conformation of human HIRA is an intrinsic feature and not caused by nucleosome binding (Supplementary Fig. 8c-d).

**Figure 4:**
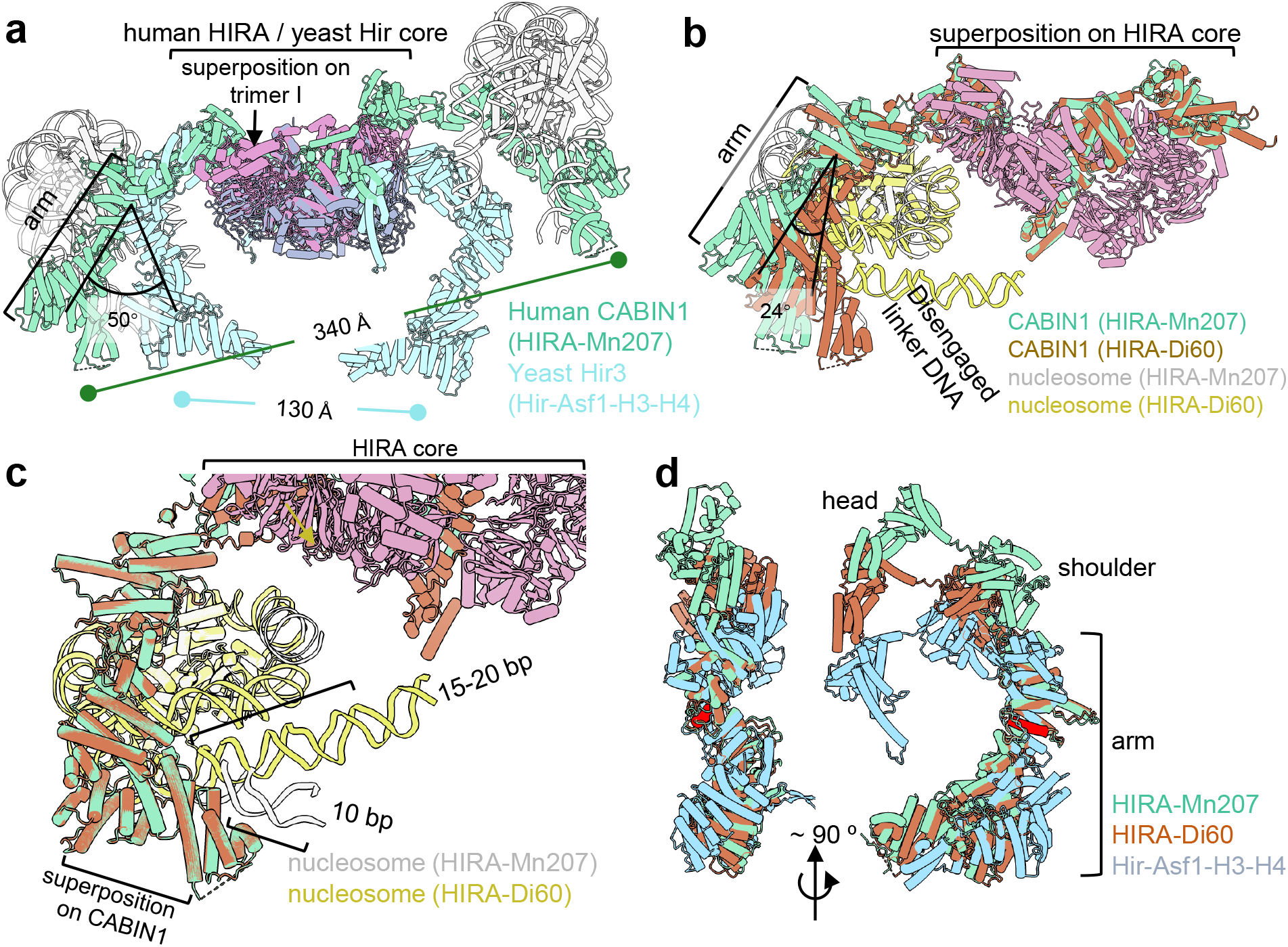
Flexible orientation of CABIN1 arms about a structurally conserved HIRA core. **(a)** Structural comparison between the human HIRA-nucleosome complex (including the second nucleosome and CABIN1 arm) and the yeast Hir-Asf1a-H3-H4 complex (PDB 8GHN). Structures were superposed using the nucleosome-proximal HIRA trimer (Trimer I) as the reference. (**b)** Structural superposition aligned on the entire HIRA core. The HIRA trimeric scaffold aligns closely between the two structures, whereas CABIN1 together with the associated nucleosome is shifted toward the HIRA core in the HIRA-Di60 (as the yellow arrow shows). **(c)** Comparison of the DNA exit region after alignment on the CABIN1 arm. In HIRA-Mn207, the CABIN1 hand interacts with and guides the exiting DNA, whereas in HIRA-Di60 the CABIN1 hand is disengaged from DNA which assumes a more curved trajectory toward HIRA core. **(d)** Internal structural differences in the yeast and human CABIN1 arm revealed by alignment on a conserved internal helix (Hir3:947-960 to CABIN1:936-949). This shows divergence of both proximal and distal segments of the CABIN1 arm, indicating flexing in the shoulder and elbow regions rather than rigid-body movement.

In the alignment of the yeast and human structures, the human CABIN1 arm adopts a ∼50° shift relative to the conserved core, while the positions of the CABIN1 head domains and shoulder are displaced only slightly (Fig. 4a; Supplementary Fig. 9a). In addition, the relative arrangement of the two HIRA trimers also differs by 12° (Supplementary Fig. 9b). To test whether the more compact yeast architecture would preclude nucleosome engagement, we docked a nucleosome onto the yeast Hir complex using the binding mode observed in human HIRA-Di60. Intriguingly, despite its smaller arm span, yeast Hir can accommodate a nucleosome in a similar manner without major steric clashes (Supplementary Fig.10, Supplementary Video 2). Together with previous evidence that yeast Hir binds DNA, this observation suggests that key features of nucleosome engagement may be conserved between yeast Hir and human HIRA^29^.

To investigate how HIRA engages chromatin in a more physiologically relevant context (with linker DNA connecting the two nucleosomes), we designed di-nucleosome substrates guided by the HIRA-Mn207 structure. There, the distance between two nucleosomes is 25-35 nm, equivalent to 60-75 bp of straight B-form DNA. Therefore, we assembled di-nucleosome substrates containing either a 60 or 75 bp linker DNA (Supplementary Fig. 11a-b). Negative-stain EM of both di-nucleosomes in complex with HIRA showed that the HIRA-Di60 sample was more homogeneous than the HIRA-Di75 complex, and it was therefore selected for further analysis (Supplementary Fig. 11c). However, although the input complex contains a fully assembled di-nucleosome, 2D class averages revealed density for only one nucleosome associated with HIRA, with only blurred density for a second nucleosome on the opposite side (Supplementary Fig. 12a, b). 3D reconstruction of the HIRA-Di60 complex revealed substantially greater structural heterogeneity compared with the HIRA-Mn207 dataset. Therefore, we rigid-body fitted the HIRA core and the nucleosome-CABIN1 module from the HIRA-Mn207 structure separately into the HIRA-Di60 density (Supplementary Fig. 13a-b).

The HIRA core adopts a similar architecture in the HIRA-Di60 and HIRA-Mn207 structures. When the two structures were aligned on the HIRA core, the CABIN1 arm and its associated nucleosome shifted toward the HIRA core in HIRA-Di60 relative to HIRA-Mn207, accompanied by an approximately 24° arm rotation, reflecting the plasticity of CABIN1 around the shoulder (Fig. 4b). We also aligned the two structures on the arm and hand region of CABIN1 and find that the CABIN1 arm clutches the nucleosome in the same manner in both structures (Fig. 4c). This indicates that CABIN1 maintains the same nucleosome-binding mode while allowing the CABIN1-nucleosome module to reposition relative to the HIRA core, possibly in response to linker-DNA constraints. Importantly, in the HIRA-Mn207 structure, ∼10 bp of straight linker DNA extending from the nucleosome is stabilized by interactions with the CABIN1 hand domain (Fig. 4c). When a dinucleosome is bound to HIRA, a longer stretch of the connecting linker DNA (∼15-20 bp) is visible at relatively high definition (Fig. 4c, Supplementary Fig. 13c for corresponding densities). However, the exiting linker DNA is dissociated from the hand domain and instead assumes a more curved trajectory towards the nucleosomal dyad and towards the HIRA core. This is likely caused by the interaction of the second nucleosome and / or linker DNA with the second CABIN1 hand domain, not resolved in our structure. Given that only UBN1 and CABIN1 directly bind DNA, UBN1 could also contribute to the redirection of DNA.

Together, the cross-species and assembly-stage comparisons identify the CABIN1 arm as the major source of HIRA adaptability, which may underlie both evolutionary tuning of HIRA-family complex size and substrate-dependent conformational changes during nucleosome assembly (Fig. 4d).

### Positioning of HIRA_s_ WD40 domains suggests a UBN1-distribution mechanism

Having established CABIN1 as the principal nucleosome and DNA-binding module of HIRA, we next focused on the HIRA_s_ WD40 domains that are visible (albeit at low definition) in our HIRA-Mn207 structure (Fig. 5a). These are connected to the HIRA trimerization domain through a ∼300 amino acid flexible linker. The HIRA_s_ WD40 domain is essential for HIRA nucleosome assembly activity and HIRA function in cells, and provides the major binding site for the UBN1 (Fig. 5b)^19,20,30–32^. Because other regions of UBN1 bind H3.3-H4 and DNA (Fig. 5b)^21^, the position of WD40 might determine UBN1 localization within the complex. We observed density for two of the six WD40 domains in our HIRA-Mn207 structure. Given the apparent two-fold organization of the complex, we infer that the corresponding WD40 domains on the opposite side could adopt similar positions but are not resolved because of the intrinsic flexibility of the extended linker connecting them to the trimerization domain (Fig. 5c, WD40 I-IV). In the yeast Hir complex, density for four WD40 domains (equivalent to WD40 II and III in our structure) are observed (Fig. 5d), while the equivalent of WD40 I is not observed in the yeast Hir-Asf1-H3-H4 structure^15^. Notably, the two WD40 domain associated with the CABIN1 arm in the yeast structure are not resolved in the HIRA-Mn207 structure, suggesting that they might be released or become highly mobile in the nucleosome-bound state^15^.

**Figure 5:**
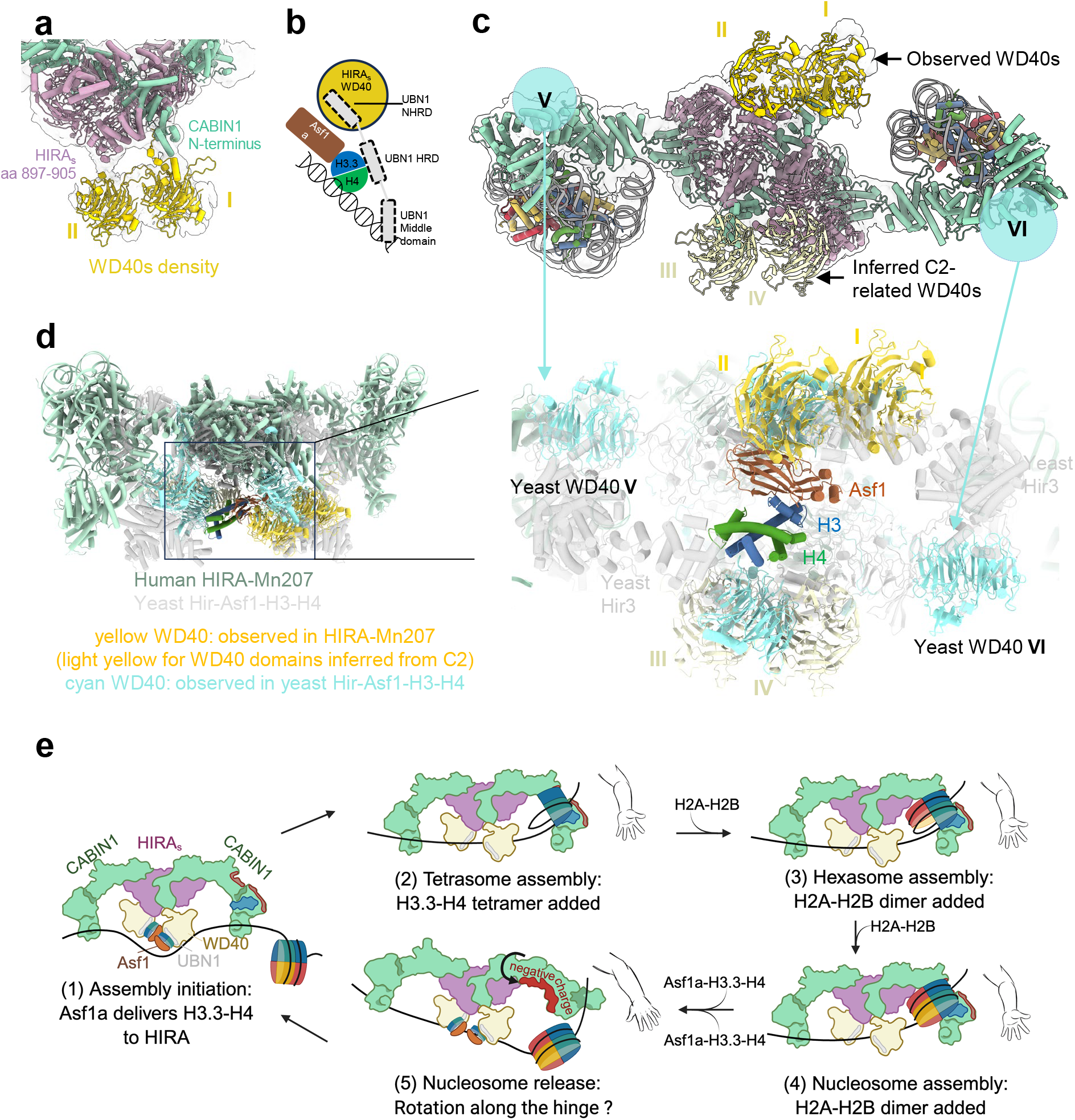
Mapping WD40 domains in the context of the entire HIRA structure, and proposed working model of HIRA-dependent nucleosome assembly. **(a)** Density assigned to WD40 domains, located near the N-terminal region of CABIN1 and HIRA_s_ (amino acids 897-905). Two AlphaFold-predicted WD40 domain models were tentatively fitted into the density. **(b)** Schematic of previously reported UBN1 interactions. UBN1 engages the HIRAs WD40 domain through its NHRD motif, binds H3.3-H4 through its HRD region and interacts with DNA through its middle domain. UBN1 itself is not resolved in the HIRA-Mn207 structure. **(c)** Overview of WD40 domain (I-IV) observed in the HIRA-Mn207 structure. WD40 V and VI that observed in yeast Hir-Asf1-H3-H4 structure were not observed in HIRA-Mn207. Roman numerals label the WD40 positions discussed in the text. **(d)** Superposition of human HIRA-Mn207 and yeast Hir-Asf1-H3-H4 (PDB 8GHN) highlights the conserved central Asf1-H3-H4-binding cavity. **(e)** Proposed working model for HIRA-dependent nucleosome assembly.

Comparison with the yeast Hir-Asf1-H3-H4 structure suggests that, despite the wider CABIN1 arm span in human HIRA, the central Asf1-H3-H4-binding region at the underside of the arch is spatially conserved between the human and yeast complexes (Fig. 5d). In yeast, two WD40 domains flank and hold the Asf1-H3-H4 substrate. This arrangement suggests that WD40-tethered UBN1 could be positioned near unassembled H3.3-H4 and DNA during assembly initiation, and such an arrangement is also compatible with the human HIRA structure. Because tetrasome formation requires two H3.3-H4 dimers, paired WD40-UBN1 modules could provide a mechanism to position two UBN1 molecules within the assembly zone, with each UBN1 engaging one H3.3-H4 dimer (initially with, then without Asf1) while its DNA-binding regions, together with CABIN1 arm and hands, helps organize the incoming DNA.

## Discussion

Our study re-defines how the massive human HIRA complex functions during replication-independent nucleosome assembly. Rather than acting as a mere H3.3 deposition factor, it appears to operate as a dynamic assembly platform that coordinates histone wrapping by DNA with a possible DNA ‘ruler’ function. We show that the complete HIRA complex (which *in vitro* does not itself discriminate between H3.3 and H3.1) contributes to all steps of nucleosome assembly in vitro and stabilizes assembly intermediates and the final nucleosome product through extensive multivalent contacts mediated primarily by CABIN1, a subunit with previously unknown function. H3.3 specificity is likely conferred through its preferential interaction with the delivery chaperone Asf1a ^15,17^.

By definition, a histone chaperone must bring DNA and histones into proximity. If promoting sequential steps in the assembly process (as we show is the case for HIRA), it should also stabilize assembly intermediates. With respect to DNA binding, we report four pronounced positively charged regions on CABIN1, two of which bind nucleosomal DNA and linker DNA, while the others might engage DNA in different ways during earlier steps in the assembly pathway. The uncharacterized UBN1 DNA binding domains (whose location in the complex are unknown) might also contribute. Histone binding is thought to be accomplished through HIRA_s_ and UBN1 subunits, based on *in vitro* binding experiments^21,33^.

The HIRAs trimerization domains form the apex of the HIRA arch. The ∼300 amino acid long disordered segment connecting to the WD40 domains provides positional flexibility to the WD40 module. Combined with the high-affinity interaction between the WD40 domain and the UBN1 NHRD motif, this flexibility may allow the WD40 domain to function as a dynamic UBN1-positioning module. Four WD40 domains could shuttle histone or DNA binding UBN1 domains at the underside of the arch to bring histones to the DNA suspended across it by the CABIN1 arms. This is consistent with the estimated presence of four UBN1 molecules in the HIRA complex^15^. Since our structures capture the endpoint of assembly, the absence of UBN1 should not be surprising as they likely function in early stages of assembly. The role of the two remaining WD40 domains (which in yeast are bound to the CABIN1 arm) remains unclear ^15,34^.

We propose a model where a stretch of >90 bp DNA is suspended across the HIRA underside via the two CABIN1 hands, where Asf1a-H3.3-H4 also bind^34^ (Fig. 5e, step 1). One of the two CABIN1 arms then promotes the wrapping of DNA around H3.3-H4 to form a tetrasome that remains bound to the arm, while the second CABIN1 arm continues to guide (or feed) the unassembled DNA (Step 2). Subsequent incorporation of two H2A-H2B dimers (which may or may not be delivered by a chaperone) generates a fully assembled nucleosome through a hexasome intermediate (Steps 3-4). This pathway is supported by our assembly assays, and by our model where CABIN1 stabilizes the product through multivalent contacts with the histone surface, nucleosomal DNA, and linker DNA. The mechanism likely requires arm movements, as suggested by our structures, allowing HIRA to remain engaged throughout the reaction rather than functioning only in the initial handoff of histones to DNA.

Our structures further suggest a potential model for how the assembled nucleosome might be released (Fig. 5e, step 5). Linker DNA trajectory can change and DNA can become disengaged from the CABIN1 hand, as seen in the HIRA-Di60 structure. During the assembly process, this redirection and disengagement could be caused by an incoming Asf1a-H3.3-H4 moiety, required for the next round of assembly. Only a slight rotation of the CABIN1 arm would then be needed to bring the extended negatively charged band on its backside into close proximity with the wrapped nucleosomal DNA, thereby facilitating nucleosome ejection, while the other hand might still guide the extended linker DNA. This speculative mechanism provides a basis for further experimentation.

From an evolutionary perspective, human HIRA exhibits higher architectural complexity than its yeast homolog, with larger subunits, more loop insertions, and a more open overall conformation. Nevertheless, it is likely that the more compact yeast complex functions through a similar mechanism, but its more narrow ‘arm-span’ might have evolved to generate the much shorter nucleosome repeat length characteristic of yeast ^35,36^. Analysis of the hand-to-hand distance suggests that yeast Hir could suspend only ∼35 base pairs of straight DNA, compared to ∼90 base pairs in human HIRA.

In metazoans, HIRA functions in the transcription-coupled deposition of H3.3 into chromatin in the wake of RNAP II^2,5,10^. AlphaFold predictions suggest a possible interface between UBN1 and a Pol II surface involving RNAP II subunits RPB11 and RPB8, with ipTM scores of 0.78 and 0.65, respectively. CABIN1 can be recruited by transcription factors, and the HIRAs WD40 domains interact with Replication Protein A (RPA), further supporting a role of HIRA in transcription^22,32^, and providing exciting new opportunities for investigation.

Many questions remain. First, what is the location and role of the UBN1 subunits? Although UBN1 is implicated in H3.3 recognition, its absence in our structure indicates substantial flexibility or only transient engagement at the end of the assembly process, the stage we have captured here. Second, many aspects of our mechanistic model presented in Fig.5e need to be confirmed experimentally, and this will require a substantial amount of sophisticated experiments. For example, although our mono-nucleosome and di-nucleosome structures suggest that changes in DNA trajectory may regulate assembly progression and product release, the extent to which the DNA path actively controls the conformation and function of the entire HIRA complex remains unknown, and it is unclear what triggers these changes. It is unknown whether HIRA functions in a processive manner, or whether it dissociates from chromatin/RNAP II during individual assembly steps. The presence of two DNA binding hands suggests the former, but this, too, remains to be tested. Third, it is unknown how the histone hand-off between Asf1a, HIRA, then DNA might function, what the role of UBN1 might be in this handoff, and how DNA wrapping is achieved. Finally, deciphering how nucleosome assembly is coupled to the transcription machinery will require a substantial effort in structural and biochemical investigation.

## Methods

### Purification of HIRA

Codon-optimized human HIRA genes (HIRA, UBN1, and CABIN1) were synthesized by GenScript. Each gene was cloned separately into pLexm vector (Addgene), with an N-terminal MBP tag on CABIN1 and an N-terminal 3×FLAG tag on UBN1. Plasmids were prepared using a Qiagen Maxiprep kit, and large-scale plasmid preparations were obtained from Azenta. Expi293F GnTI^−^ cells (Thermo Scientific) were cultured in FreeStyle 293 Expression Medium (Thermo Scientific) supplemented with 1% FBS. For transfection, a total of 1 mg plasmid DNA (0.35 mg per construct) was mixed with 1 mg PEI (Polysciences) and added to 1 L of cells. Cells were harvested after 60-72 hours of expression.

Purification of the HIRA complex was performed in three steps: (1) FLAG affinity purification, (2) heparin chromatography to remove nucleic acid contamination, and (3) MonoQ ion exchange chromatography to improve complex homogeneity. For lysis, 50 mL of lysis buffer (20 mM HEPES pH 8.0, 500 mM NaCl, 0.5 mM TCEP, 5% glycerol, and EDTA-free protease inhibitor cocktail) was added to cells from 1 L culture. Cells were lysed by sonication for 5 minutes at 50% power (3 s on, 8 s off). The lysate was clarified by centrifugation at 18,000 rpm for 1 hour, and the supernatant was incubated with Anti-DYKDDDDK G1 affinity resin (GenScript) for 2 hours at 4°C. After washing with 20 column volumes of lysis buffer, the HIRA complex was eluted by incubation with 0.5 mg/mL FLAG peptide (DYKDHDGDYKDHDIDYKDDDDK, ABclonal Science) for 1 hour. The eluted sample was then loaded onto a heparin column equilibrated in buffer A (20 mM HEPES pH 8.0, 100 mM NaCl, 0.5 mM TCEP, 5% glycerol) and eluted with a linear gradient to buffer B (20 mM HEPES pH 8.0, 1.5 M NaCl, 0.5 mM TCEP, 5% glycerol) (Supplementary Fig. 14a). The major peak fractions were pooled and diluted to 100 mM NaCl before loading onto a MonoQ column using the same buffer system. Due to the heterogeneous stoichiometry of the HIRA complex, the MonoQ elution profile was broad. Fractions were therefore selected based on SDS-PAGE analysis showing equal amounts of the three subunits (Supplementary Fig. 14b). Target fractions were concentrated to ∼1 mg/mL and exchanged into storage buffer (20 mM HEPES pH 8.0, 250 mM NaCl, 0.5 mM TCEP, 5% glycerol). Protein samples were aliquoted, flash-frozen in liquid nitrogen, and stored at -80°C. Typically, ∼0.2 mg of purified HIRA complex was obtained from 1 L of mammalian cell culture.

### Nucleosome assembly intermediates

Human core histones were obtained from The Histone Source (https://histonesource-colostate.nbsstore.net/). The DNA sequence used for reconstitution is shown below, with the Widom 601 positioning sequence underlined. DNA fragments were ordered from IDT and amplified using OneTaq 2× Master Mix (NEB).

#### 207bp DNA (for nucleosome)

ATCTAATACTAGGACCCTATACGCGGCCGC<u>ATCGGAGAATCCCGGTGCCGAGGCCGCTCAA TTGGTCGTAGACAGCTCTAGCACCGCTTAAACGCACGTACGC</u>**<u>G</u>**<u>CTGTCCCCCGCGTTTTAA CCGCCAAGGGGATTACTCCCTAGTCTCCAGGCACGTGTCAGATATATACATCGAT</u>TGCATGT GGATCCGAATTCATATTAATGAT

#### 128bp DNA (for Hexasome)

CCTATACGCGGCCGCATCGGAGAATCCCGGTGCCGAGGCCGCTCAATTGG<u>TCGTAGACAG CTCTAGCACCGCTTAAACGCACGTACGCGCTGTCCCCCGCGTTTTAACCGCCAAGGGGATTACTCCCT</u>

#### 79bp DNA (for tetrasome)

<u>TCGTAGACAGCTCTAGCACCGCTTAAACGCACGTACGCGCTGTCCCCCGCGTTTTAACCGCCAAGGGGATTACTCCCTA</u>

#### 354 bp DNA (for dinucleosome Di60

<u>GAGGCCTAACGAGCGCCTTGGTATCCGGAACGGGTTCTAGTACCCGTTAGCGTGGTTTAGAGGGGCAAAGGAACATCTTTCCCCCCCCCGAGATACGGGCACCGTTCGGACCCTGGTTAGT CCAGTGCTACTGCCGGTTCCTAGCCC</u>TTCGAGCTCTCACAGATCTGAACTAGTATTAAGCGT ACTGGATCCAAGTAAAGGTCGCGT<u>AAGCCTTGTCGCTCTGCGATTCGATGAGAGTCGACCG GTGCCGGGTATTGTATCGCCGCTTGACTCTGGAATGGCGTCTACGTAGCGCTAAGCACTATTAGAGTCCTCTCCTGCATCTCCAAGGATAGTGCCTATAAATCGTCCACC</u>

#### 369 bp DNA (for dinucleosome Di75

<u>GAGGCCTAACGAGCGCCTTGGTATCCGGAACGGGTTCTAGTACCCGTTAGCGTGGTTTAGAGGGGCAAAGGAACATCTTTCCCCCCCCCGAGATACGGGCACCGTTCGGACCCTGGTTAGT CCAGTGCTACTGCCGGTTCCTAGCCC</u>TTCGAGCTCTCACAGATCTGAACTAGTATTAAGCGT ACTGGATCCAAGTAAAGGTCGCGTCGACCTAATGACGGA<u>AAGCCTTGTCGCTCTGCGATTC GATGAGAGTCGACCGGTGCCGGGTATTGTATCGCCGCTTGACTCTGGAATGGCGTCTACGTAGCGCTAAGCACTATTAGAGTCCTCTCCTGCATCTCCAAGGATAGTGCCTATAAATCGTCCA CC</u>

### Fluorescent labeling of histones

Human histone mutants containing single cysteine substitutions, H2B T115C and H4 E63C, were used for site-specific fluorescent labeling^37^. Lyophilized histones were dissolved in unfolding buffer containing 7 M guanidine hydrochloride, 20 mM NaCl, 20 mM Tris-HCl pH 7.5, and 0.1 mM TCEP, and incubated for 1 h at room temperature to fully solubilize the proteins. Protein concentration was determined by absorbance at 280 nm. A 1:1 molar ratio of maleimide-conjugated fluorophore was then added to the unfolded histones. H2B T115C was labeled with ATTO 647N maleimide (Invitrogen), and H4 E63C was labeled with Alexa Fluor 488 maleimide (Invitrogen). Labeling reactions were carried out overnight at 4 °C with gentle rotation. Labeled histones were then mixed with the corresponding unlabeled histones at the desired stoichiometry and adjusted to a final protein concentration of approximately 1 mg/ml. Histone complexes were refolded by dialysis against refolding buffer (20 mM NaCl, 20 mM Tris-HCl pH 7.5, 0.1 mM TCEP and 0.5mM EDTA) overnight at 4 °C, with two additional buffer exchanges. Refolded histone complexes were concentrated and further purified by size-exclusion chromatography using a Superdex 200 column. Labeling efficiency was estimated by absorbance spectroscopy based on the concentration of fluorescent dye relative to the concentration of refolded histone complex.

### Nucleosome reconstruction

Nucleosome, hexasome and tetrasome reconstitution were performed according to previously published protocols^38,39^ and analyzed by native PAGE. Di-nucleosomes were reconstituted similarly but with twice the amount of histone added. A BamHI restriction site was inserted into the linker region of DNA used for di-nucleosome reconstitution. We combined BamHI digestion and mass photometry to assess the quality of the di-nucleosome.

### Nucleosome assembly assay

The nucleosome assembly assay was performed as described^27^. 147bp Widom 601 DNA was used, sequences are given above. The reaction buffer contains 20 mM HEPES pH 7.5, 150 mM NaCl, 5% glycerol, 0.5 mM TCEP. For tetrasome assembly, 100 nM DNA and 150 nM H3.3-H4 tetramer was incubated with 20-40 nM recombinant HIRA for 15 min at room temperature. For nucleosome assembly, 200 nM H2A-H2B dimer was added in addition (Fig. 1d). Because HIRA binds the assembled product, when more than 40 nM HIRA was included in the reaction, the product became bound by HIRA and remained in the well due to its large size. In the H2A-H2B deposition assay, 500 nM competitor DNA was added to release assembled intermediates (Supplementary Fig. 1a-1b).

### Fluorescence polarization binding assay

To determine the binding affinity between HIRA and various nucleosome components, we mixed varying amounts of HIRA with 5 nM Atto-647-labeled H2A-H2B dimers and Alexa-488-labeled H3.3/H3.1-H4 tetramers, as well as 5 nM 147 bp nucleosomes labeled with Alexa-488 on H4. The binding buffer was 20 mM HEPES (pH 7.5), 150 mM NaCl, 5% glycerol, 0.01% NP-40, 0.01% CHAPS, and 0.5 mM TCEP. Reactions were incubated in a 384-well microplate for 10 min. at room temperature. Fluorescence polarization data were acquired using a CLARIOstar microplate reader (BMG Labtech) and analyzed with GraphPad Prism. The binding data were fitted using a one-site binding model:

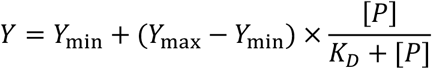

where *Y* is the measured polarization, *Y*_min_ and *Y*_max_ represent the minimum and maximum polarization values, respectively, [*P*] is the protein concentration, and *K*_*D*_ is the apparent dissociation constant. Three replicates were performed, and measurements obtained from the same batch were reproducible.

### EM sample and grid preparation

For HIRA-Mn207 and HIRA-Di60 samples: We mixed 90 pmol HIRA (assume 1mg/ml HIRA equal to 1pmol HIRA) with 108 pmol 207bp nucleosome or 55 pmol di-nucleosome. GraFix was performed to stabilize the complex^40^. The GraFix buffer consisted of 20 mM HEPES pH 7.5, 50 mM NaCl, with 15% sucrose in buffer A and 45% sucrose, 0.04% glutaraldehyde in buffer B. The sample was centrifuged using an SW55Ti rotor at 22,500 rpm for 14 h at 4 °C. Density gradients and fractionation were conduct using the Gradient Station and Piston Gradient Fractionator from BioComp Instruments. Fractions were quenched with 100 mM Tris pH 7.5, combined, and dialyzed overnight against freezing buffer (20 mM HEPES pH 7.5, 50 mM NaCl, 2.5% glycerol, and 0.5 mM TCEP). The samples were then concentrated for grid preparation.

For HIRA-Asf1a-H3.3-H4 complex, we mixed 100 pmol HIRA with 250 pmol Alexa-488-labeled H3.3-H4 tetramer and 250 pmol Asf1a (purchased from The Histone Source, Fort Collins, CO). GraFix was performed to stabilize the complex as above. The GraFix buffer consisted of 20 mM HEPES pH 7.5, 150 mM NaCl, with 15% sucrose in buffer A and 45% sucrose, 0.04% glutaraldehyde in buffer B. Sample centrifugation and fractionation procedures are same as HIRA-Mn samples.

### Negative-stain EM

A 400-mesh copper grid (Electron Microscopy Sciences) with a continuous carbon-supporting layer was used to prepare negative-staining samples. 100 nM crosslinked HIRA complex was applied to glow discharged grids and incubated for 30 s. After incubation, the grids were stained by five successive 35ul drops of 2% (w/v) uranyl formate solutions for 3s each and then blotted to dryness. Data were collected using an FEI Tecnai F30 transmission electron microscope (TEM). Data processing was performed with cryoSPARC^41^ to create a reference map for subsequent cryo EM data processing.

### Grid preparation of crosslinked HIRA-Mn and HIRA-Di60 complex

0.045% β-OG was added to a 150-200 nM sample immediately prior to freezing to improve particle distribution and ice thickness. A thin carbon layer (∼1.5 nm) was floated onto Quantifoil R1.2/1.3 Au 300-mesh grids (Electron Microscopy Sciences) using the Leica ACE 600 carbon evaporator. Then 4 µL sample was applied to gently glow-discharged, carbon-coated grids and vitrified using a Vitrobot Mark IV (Thermo Fisher). Blotting was performed for 2 s at 100% humidity.

### Cryo-EM data collection and processing

Images were collected at nominal magnification of 130,000 × (Pixel size: 0.91 Å) on a FEI Titan Krios (300 kV), equipped with a Falcon 4 Direct Detection Camera. The movies were captured in super resolution mode with a total dose of 50 e/Å^2^ (electrons per Angstrom square). Defocus range was −0.6 to −2.5 μm.

The HIRA-207bp nucleosome dataset was pre-processed (motion correction and CTF estimation) and processed with CryoSPARC(v4)^41,42^. Particle picking was performed using a combination of manual picking and Topaz training^43^. The particle diameter was set to 400 Å, and particles were extracted with a box size of 768 pixels and binned 4× for 2D classification. Classes displaying nucleosome features and those showing the HIRA scaffold were selected separately for ab initio reconstruction to generate independent initial reference maps. All classes were subsequently subjected to heterogeneous refinement to remove low-quality particles. Multiple rounds of ab initio reconstruction and heterogeneous refinement were performed to further clean the dataset. The selected particles were then re-extracted without binning and subjected to non-uniform refinement to obtain the highest-resolution map^44^. For the CABIN1-nucleosome map (only including CABIN1 C-terminal nucleosome binding motif), masks were created in Chimer X^45^ to mask on overall CABIN1-nucleosome and CABIN1 alone separately for subsequent local refinement. Then 3D classification with different masks was performed in CryoSPARC to class out bad particles and distinct conformational states. The best classes were selected for final refinement. The final resolution of CABIN1-nucleosome is 3.38Å, CABIN1 alone is 6.95Å. Similar strategy was applied to HIRA core processing. The final resolution of HIRA core without WD40 domain is 3.69Å, with WD40 domain is 3.94Å (Supplementary Fig.15 and 17).

3D reconstruction of the HIRA-Di60 complex revealed substantially greater structural heterogeneity compared with the HIRA-Mn207 dataset. As above, we combined RELION and CryoSparc to solve the HIRA core and nucleosome-CABIN1 part separately. We were unable to obtain a high-resolution reconstruction of the entire HIRA core for the HIRA-Di60 dataset. Instead, we reconstructed a low-resolution map in RELION that shows density for the entire HIRA core, as well as partial density for the nucleosome and its associated CABIN1 arm. By selecting 2D classes that displayed clear nucleosome features, we were able to resolve the nucleosome-CABIN1 region to 3.8-7.2 Å, comparable the resolution obtained in the HIRA-Mn207 dataset. This was sufficient to support reliable modeling of the nucleosome and the interacting CABIN1 subunit, while the definition of the core was only sufficient for rigid body fitting. To interpret the overall architecture of the HIRA-di-nucleosome complex, we opted for a rigid-body fit of the HIRA-Mn207 model into the low-resolution map (Supplementary Fig. 16-17).

### Cross-linking and mass spectrometry

HIRA (0.5 μM) was incubated with 147 bp nucleosome at a 1:1 molar ratio in crosslinking buffer (20 mM HEPES pH 7.5, 50 mM NaCl, 0.5 mM TCEP). The mixture was incubated on ice for 10 min, followed by the addition of DSSO crosslinker (Sigma) to a final concentration of 2 mM, and further incubated on ice for 2 h. 5 μL sample was used for analysis by by SDS-PAGE to assess crosslinking efficiency, and the remaining sample was flash-frozen and submitted to the UConn Proteomics & Metabolomics Facility for mass spectrometry analysis. The interactions were confirmed by three reproducible replicates. The CX-Circos tool was used to visualize intra- and intermolecular interactions within HIRA and between HIRA and the nucleosome (https://cx-circos.vercel.app/app).

### Model building

Initial model building of the HIRA core was guided by the structure of yeast Hir, which forms a dimer through two heterotrimers (two Hir1 and one Hir2 molecules) that are bridged by the Hir3 head and N-terminal arm. We used AlphaFold3 to predict a partial model of the corresponding much larger human core complex, with an input of six HIRAs (amino acids 680-1017) and two CABIN1 (amino acids 1-686). The predicted structure shows a similar overall organization as the yeast Hir core with relatively high confidence, despite low sequence similarity (∼20-25%; Supplementary Fig.18). This model was used for HIRA core model building.

For the nucleosome, the PDB 1AOI^46^ model was fitted into the density, and the linker DNA was extended and adjusted in ISOLDE^47^. For the CABIN1 region, we use AlphaFold to predict individual regions separately, excluding flexible loops, and then fitted these models into the density. The final model was completed through iterative refinement in Coot^48^ and Phenix^49^.

## Author contributions

W.T. performed all aspects of this work, with help from co-authors as indicated: L.Y and V.K helped with optimization of the carbon-covered grid and grid preparation procedure. S.C and V.K. helped and advised with data processing. K.L. assisted with interpretation of data, writing, and figure preparation.

## Acknowledgements

Supported by the Howard Hughes Medical Institute (WT and KL). S.C. is a JCC-HHMI fellow funded by the Jane Coffin Childs Memorial Fund for Medical Research. Cryo-EM imaging was assisted by Dr. Erik Hartwick, and data collection was performed at the Biochemistry Krios Electron Microscopy Facility (BioKEM) at CU Boulder (RRID: SCR_019057). A portion of this research was supported by NIH grant R24GM154185 and performed at the Pacific Northwest Center for Cryo-EM (PNCC) with assistance from Rose Marie. We thank Rakshaa Mureli for help with cell culture and HIRA expression, and Dr. Liqi Yao for help with optimizing carbon grids. We thank Dr.Siyu Chen and Dr. Vignesh Kasinath for help with data processing. We acknowledge Dr. Yiyan Yang for discussion and help generate the working model. We acknowledge Dr. Annette Erbse and the Shared Instruments Pool (RRID: SCR_018986) of the Department of Biochemistry at the University of Colorado Boulder for providing access to the Avanti JXN 26 Super Speed Centrifuges and rotors, funded by NIH Grant (R24OD033699-01). Software used in this project was curated by SBGrid. This work utilized the Blanca condo computing resource at the University of Colorado Boulder. Blanca is jointly funded by computing users and the University of Colorado Boulder. Data storage supported by the University of Colorado Boulder PetaLibrary.

## Data availability

The cryo-EM micrographs, cryo-EM maps and model coordinates have been made available on Electron Microscopy Public Image Archive (EMPIAR), Electron Microscopy Data Bank (EMDB) and Protein Data Bank (PDB), respectively. HIRA-Mn207 complex: EMD-77049; EMD-77044: EMD-77039; EMD-77041; EMD-77040; EMD-77050 and PDB: 13FQ. HIRA-Di60 complex: EMD-77078; EMD-77089; EMD-77076; EMD-77077 and PDB: 13IA.

