## supplementary figures for "Structure of the human HIRA histone chaperone with a nucleosome suggests a stepwise nucleosome assembly mechanism"

### Supplementary Figures 1-18

### Supplementary Video 1

Overview of the entire HIRA-nucleosome complex based on C2 symmetry.

### Supplementary Video 2

Model showing that the yeast Hir complex can accommodate nucleosomes. Superposition of HIRA-Di60 CABIN1-nucleosome binding module with the yeast Hir-Asf1-H3-H4 structure (PDB 8GHN). The structures were aligned based on the CABIN1 upper arm, using CABIN1 residues 936-949 and Hir3 residues 947-960. Human HIRA components were removed to generate the yeast hypothetical “Hir-nucleosome” model.

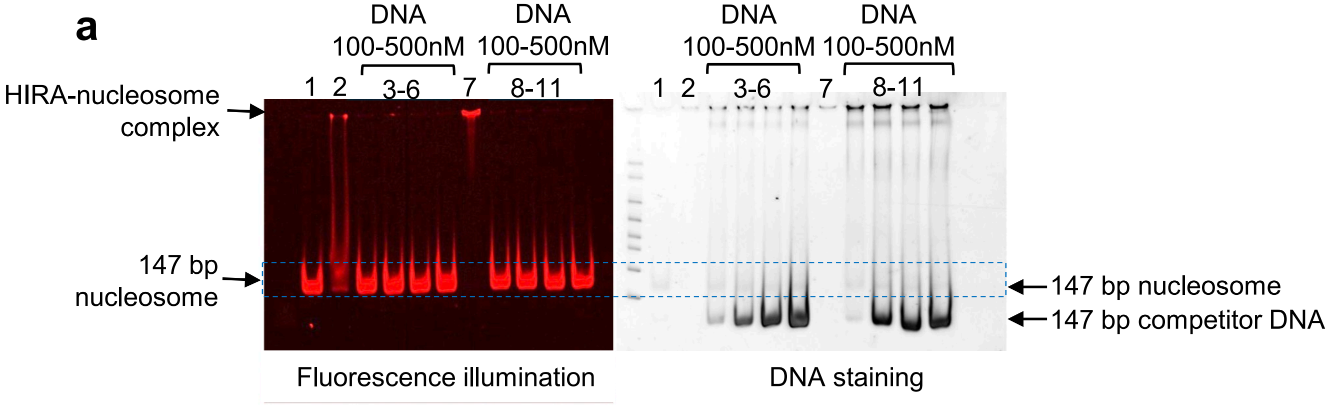


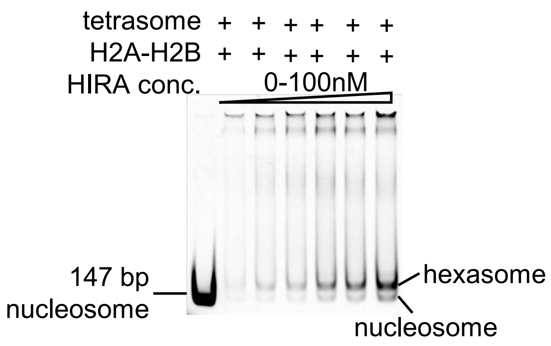


**Supplementary Figure 1: HIRA-mediated nucleosome assembly. (a) DNA competes nucleosome from HIRA.** Because HIRA binds tightly to its assembly product, exogenous DNA was added to release HIRA following nucleosome assembly. Samples were analyzed by 5% native PAGE. Lane 1: 100 nM 147 bp nucleosome control, with Alexa-488 labeled histone H4. Lanes 2 and 7: 100 nM nucleosome was incubated with HIRA at a 1:1 or 1:2 molar ratio. Without competitor DNA, HIRA and nucleosomes form a complex that is trapped in the well due to its large size. Lanes 3-6 and 8-11: addition of increasing concentrations of competitor DNA (100–500 nM) to the reactions in lanes 2 and 7, respectively. Efficient release of nucleosomes upon addition of competitor DNA indicates that DNA can compete nucleosome products away from HIRA. Based on this result, 500 nM competitor DNA was used in subsequent HIRA-dependent assembly assays. The left panel shows fluorescence imaging, selectively visualizing nucleosomes and associated species via 488-labeled H4. The right panel shows DNA staining, revealing both free DNA and nucleosome bands. The blue dashed box indicates the position of nucleosome bands detected in both fluorescence and DNA-stained images. (**b) HIRA promotes the incorporation of H2A-H2B into tetrasomes to form nucleosomes**. 100 nM H2A-H2B dimer was incubated with varying amounts of HIRA and then 100 nM tetrasome was added to initiate nucleosome assembly. At low HIRA concentrations, the primary product is hexasome, while increasing HIRA levels lead to the formation of more nucleosomes. Assembly products were analyzed by native PAGE and visualized by EtBr.

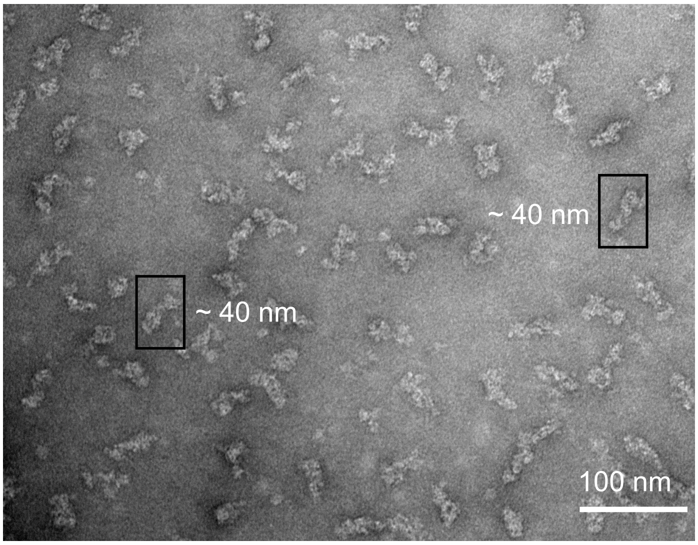


**Supplementary Figure 2: Negative-stain EM evaluation of the HIRA-Mn207 complex.** Negative-stain EM of crosslinked HIRA-Mn207 complexes shows homogeneous particles with an average length of ~ 40 nm. Black boxes indicate representative particles.

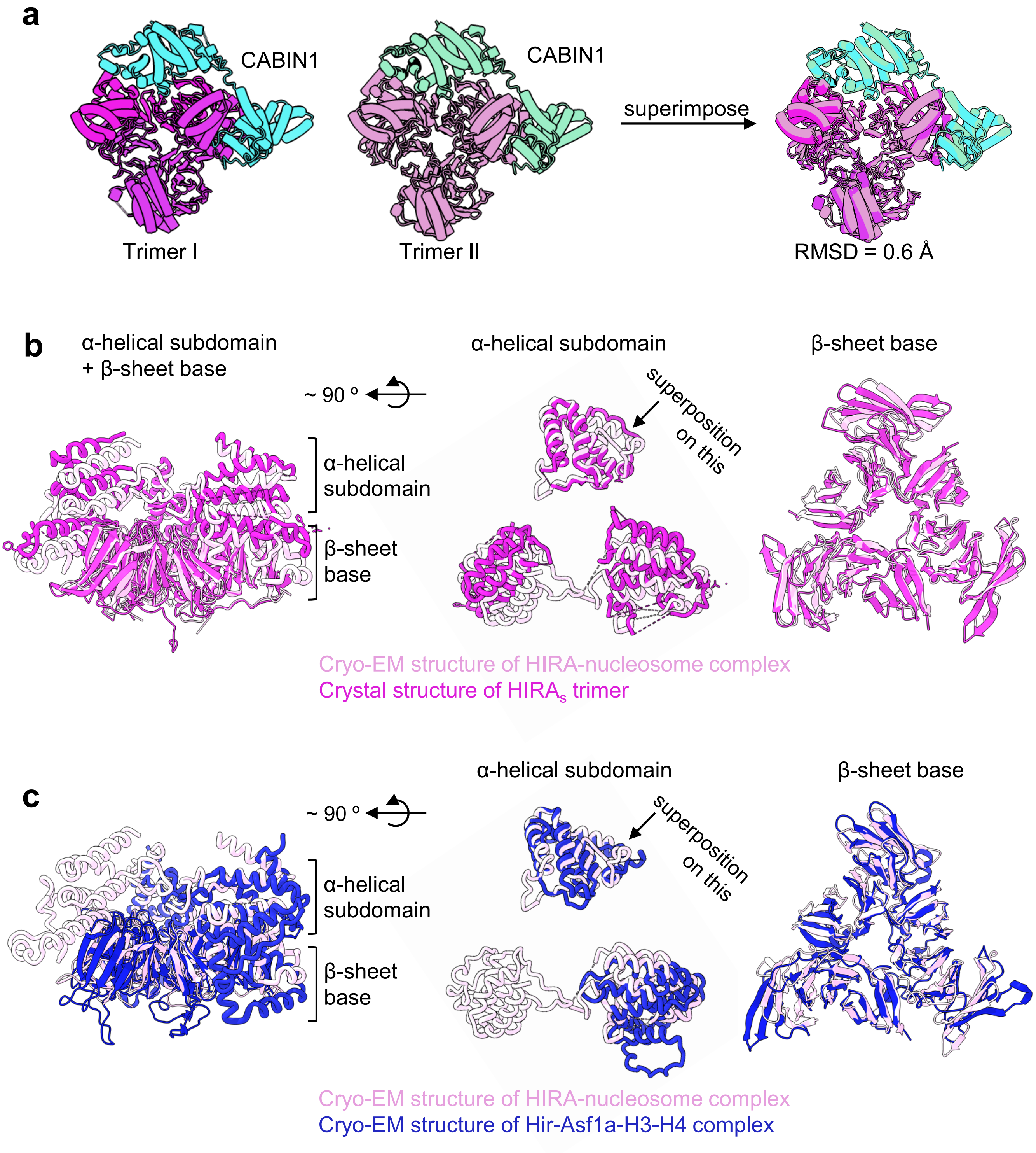


**Supplementary Figure 3: Structural comparison of the central HIRA$_s$ trimer in human and yeast. (a)** Superposition of the two HIRA trimers together with their associated CABIN1 head and shoulder domains. **(b)** Superposition of the HIRA in our structure (light pink) with the crystal structure of the human apo HIRA trimer (PDB 5YJE, magenta). Views highlighting the combined α-helical subdomain and β-sheet base (left), the isolated α-helical subdomains upon 90° rotation (middle), and the β-sheet base (right) are shown. The comparison reveals strong conservation of the central β-sheet base, with greater variability in the relative positions of the α-helical subdomains. **(c)** Superposition of the HIRA trimer from our structure (light pink) with the corresponding trimeric core from the yeast Hir-Asf1-H3-H4 cryo-EM structure (PDB 8GHN, blue). Orientations are the same as in a). The central β-sheet base is well conserved between the human and yeast complexes, whereas the α-helical subdomains adopt distinct orientations.

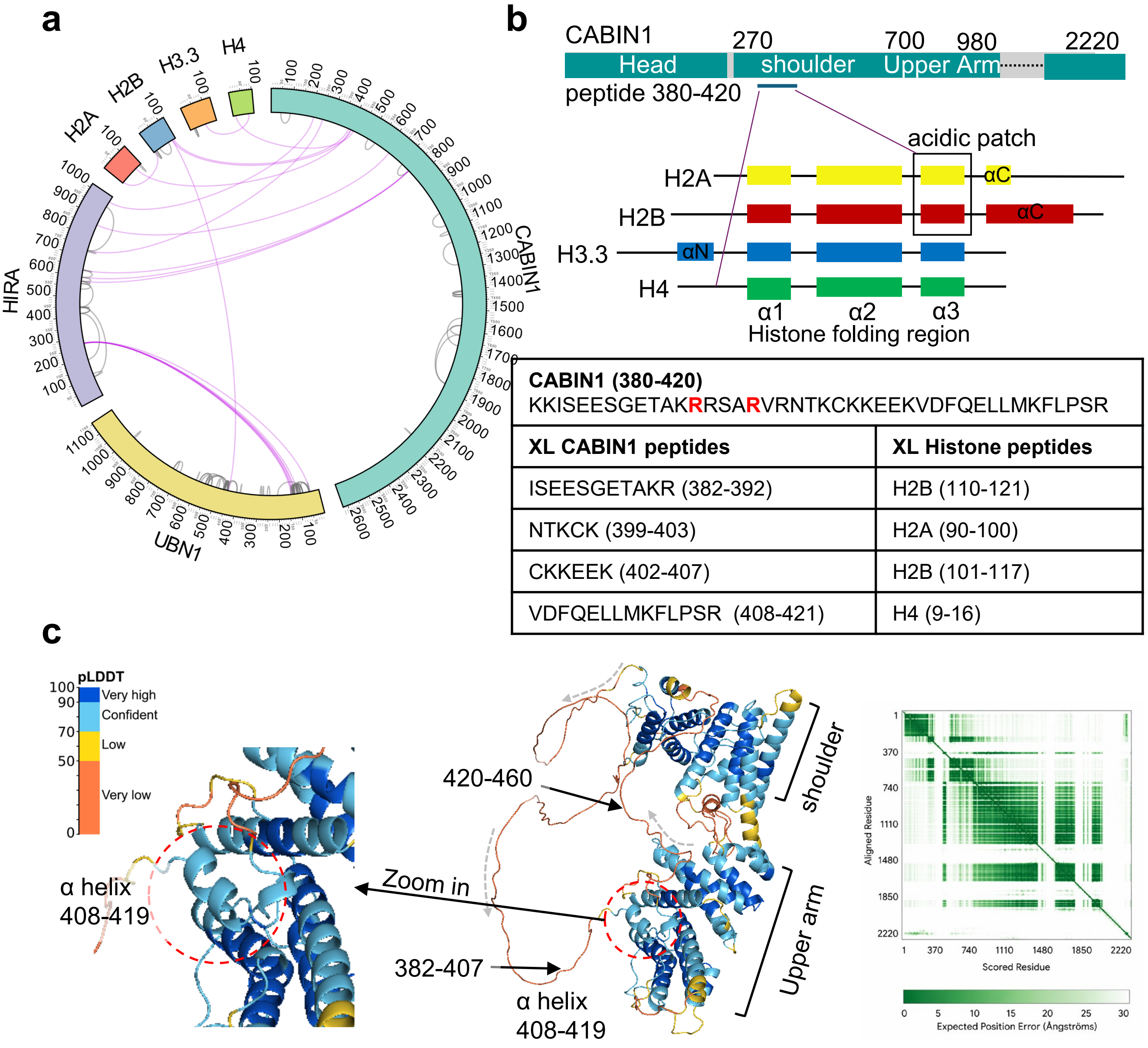


| CABIN1 (380-420) KKISEESGETAKRRSARVRNTKCKKEEKVDFQELLMKFLPSR | |
|---|---|
| **XL CABIN1 peptides** | **XL Histone peptides** |
| ISEESGETAKR (382-392) | H2B (110-121) |
| NTKCK (399-403) | H2A (90-100) |
| CKKEEK (402-407) | H2B (101-117) |
| VDFQELLMKFLPSR (408-421) | H4 (9-16) |

**Supplementary Figure 4. Identification of a CABIN1 peptide that interacts with the nucleosomal acidic patch. (a)** Crosslinking mass spectrometry (XL-MS) analysis showing that CABIN1 is the main HIRA subunit that interacts with nucleosomal histones. **(b)** Mapping XL-MS interactions onto domain diagram of CABIN1 and histones. Crosslinks localize a CABIN1 peptide (amino acids 380-420) to the nucleosomal acidic patch formed by H2A-H2B (amino acid 95-105 and 125-145) , and to the N-terminal tail of H4. The table shows all the crosslinked peptides identified between CABIN1 and histones. The two arginine anchors are highlighted in red. **(c)** AlphaFold prediction of full length CABIN1 highlighting the region encompassing residues 382-460. The model suggests that an α-helix (amino acid 408-419) and adjacent loops extrudes from the shoulder domain toward the upper arm domain. It packs into the interface formed by two helices in the upper arm (amino acids 813-832 and 887-901) and thereby positions the adjacent acidic-patch-binding loop (RRSAR peptide) containing the arginine anchor (see Fig. 3a-b)

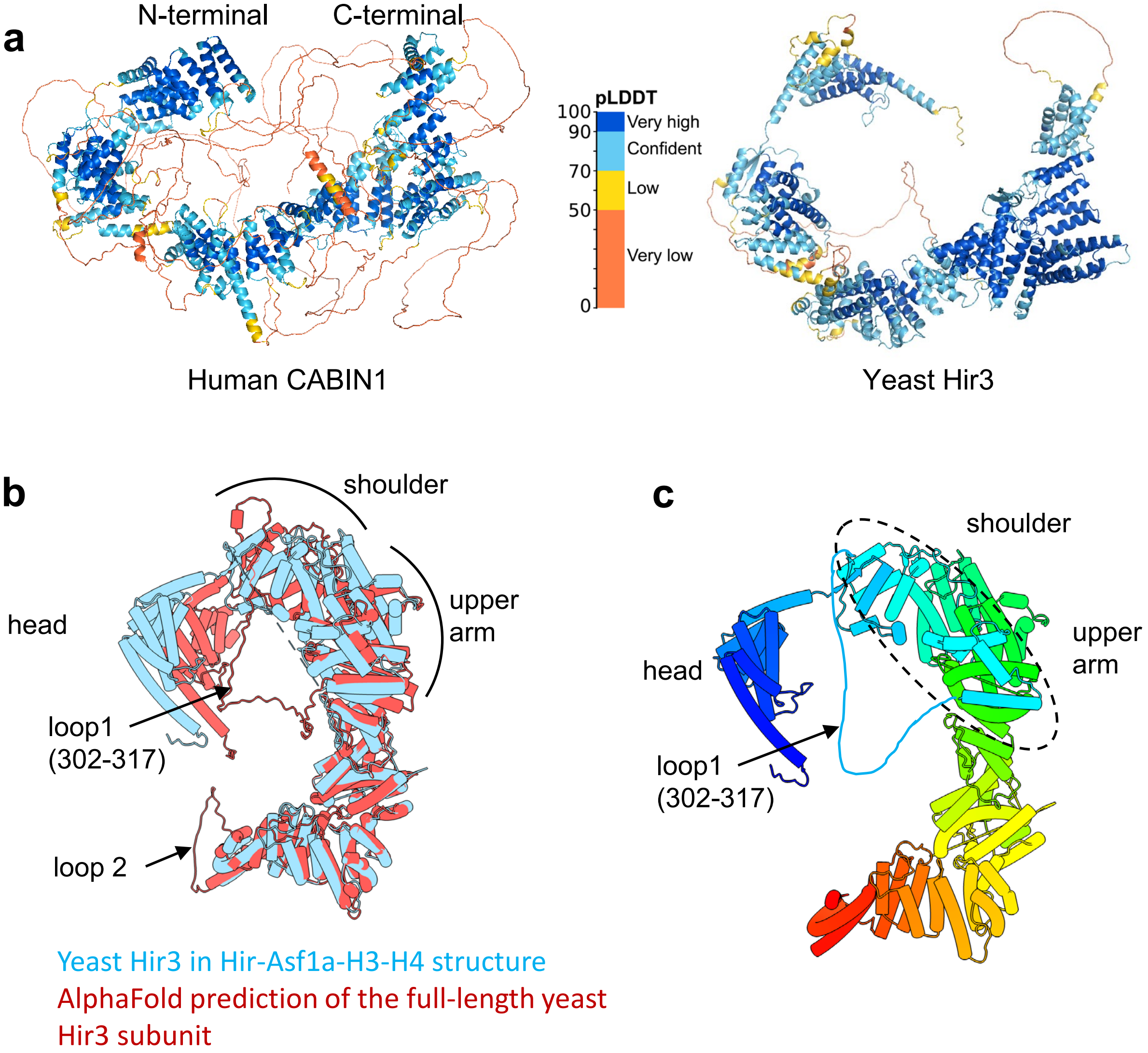


**Supplementary Figure 5. AlphaFold prediction of full length human CABIN1 and yeast Hir3. (a)** The AlphaFold predicted human CABIN1 and yeast Hir3 structures are similar, except for in loop regions. **(b)** Structural superimposition of yeast Hir3 from the Hir-Asf1a-H3-H4 structure (PDB 8GHN) and the AlphaFold-predicted full-length yeast Hir3. The predicted full-length Hir3 exhibits two loops. Loop 1 (amino acids 302-317) extends from the shoulder domain to the upper arm; it contains 8 arginine and 7 lysine residues (R/K = 20.3%), suggesting a conserved capacity to recognize the acidic patch. **(c)** Structure of yeast Hir3 shown in a rainbow color scheme, with loop1 manually traced. In yeast, unlike in human HIRA, loop1 does not rely on an α-helix for anchoring. Instead, an α-helical bundle from the shoulder region (indicated by dotted lines) extends toward the upper arm, positioning the loop in a similar spatial arrangement.

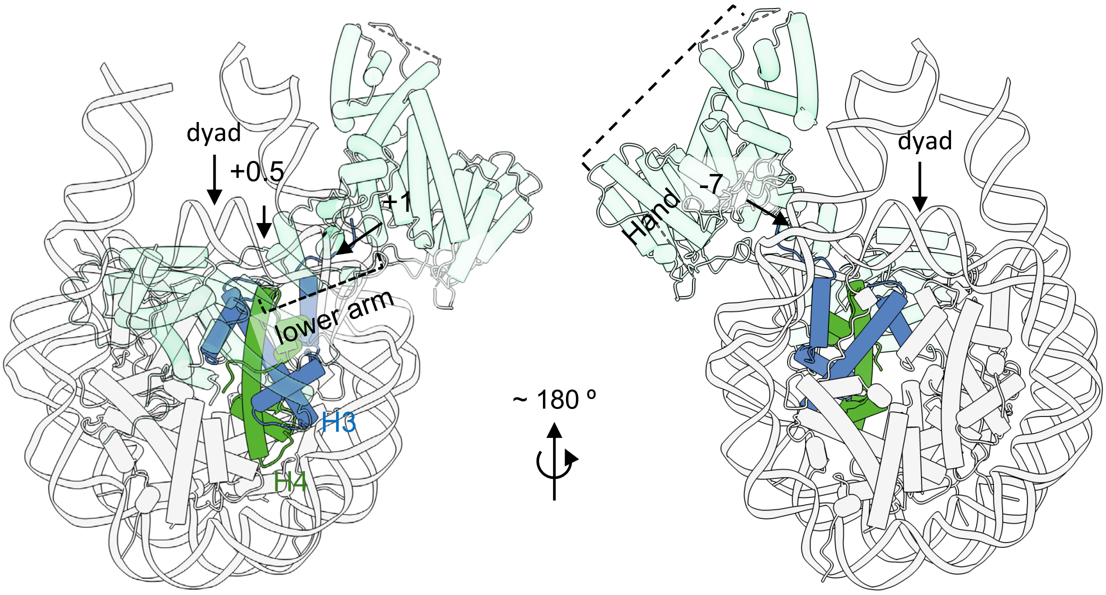


**Supplementary Figure 6: Positioning of the CABIN1 subunit relative to nucleosomal DNA superhelical locations (SHLs).** CABIN1 tracks along the DNA gyre spanning approximately SHL +0.5 to +1 and SHL −7.

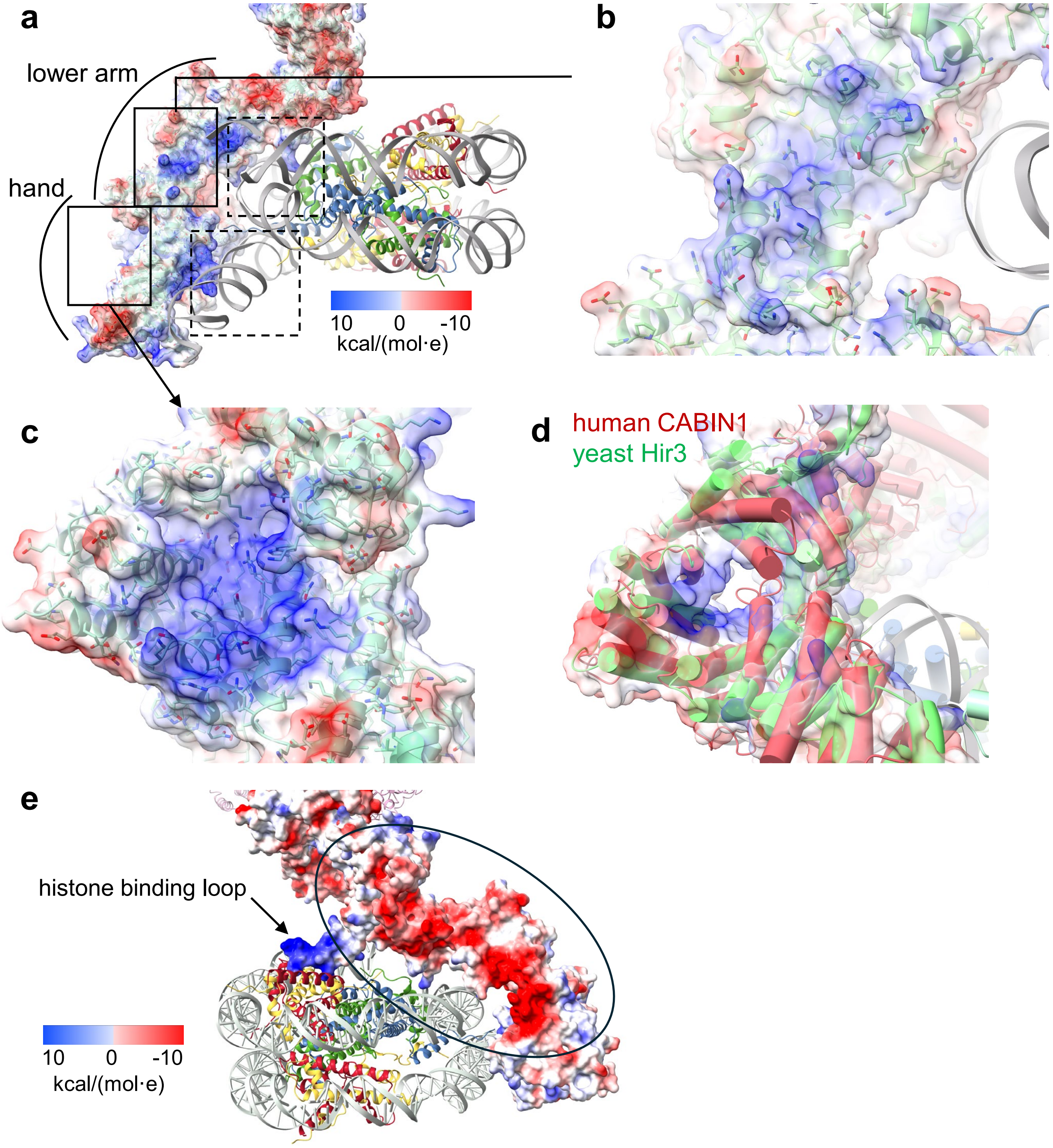


**Supplementary Figure 7 : Charge distribution on the CABIN1 arm. (a)** Electrostatic surface representation of the CABIN1 arm highlighting multiple positively charged patches. Dashed boxes mark the two DNA-binding pockets discussed in the main text that interact with nucleosomal DNA, whereas solid boxes indicate additional basic patches of currently unknown function (enlarged in b, c, d). **(b)** Basic surface located along the lower arm of CABIN1 positioned near the nucleosomal DNA gyre. **(c)** Additional positively charged patch on the back side of the nucleosomal DNA-binding surfaces close to the distal hand. (**d**) The 'basic channel' in yeast Hir3 (blue), hypothesized to bind ss DNA or RNA is shown as electrostatic surface. This channel is closed in our structure of the human HIRA-nucleosome complex, due to the shift of two helices toward the center of the channel, partially occluding the opening. **(e)** A pronounced negatively charged band along the backside of the upper and lower arm of CABIN1 (indicated by oval), of currently unknown function. Arrow indicates the  histone binding loop (RRSAR peptide) reaching towards the acidic patch of the nucleosome (see Fig. 3).

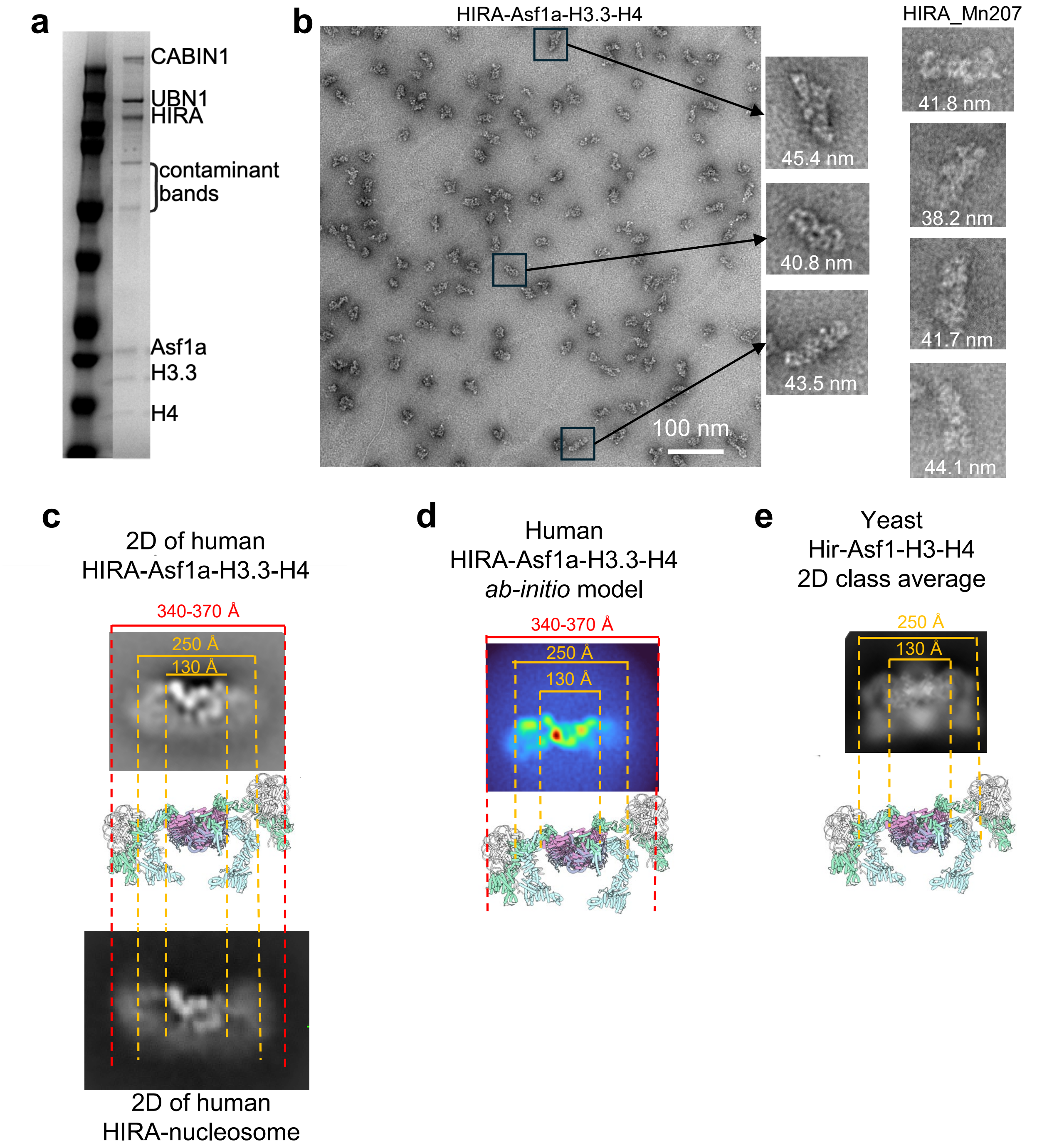


**Supplementary Fig. 8: Estimated dimensions of the human HIRA-Asf1a-H3.3-H4 complex. (a)** Asf1a co-elutes with the HIRA complex and H3.3-H4 substrate from a MonoQ column. **(b)** Negative-stain EM evaluation of the HIRA-Asf1a-H3.3-H4 complex and HIRA-Mn207 complex using particles from Supplementary Fig. 2. Apparent particle maximum dimensions were measured manually from isolated particles in Fiji after scale-bar calibration. **(c)** The human HIRA-nucleosome structure and the yeast Hir-Asf1-H3-H4 structure were mapped onto 2D class averages obtained with the human HIRA-Asf1a-H3.3-H4 complex and the human HIRA-nucleosome complex, to estimate the overall dimensions of the human HIRA-Asf1a-H3.3-H4 complex. Dashed outlines indicate reference dimensions corresponding to structural features shown in Fig. 4a: 130 Å marks the distance between the two yeast Hir3 hand regions, 250 Å marks the overall width of the yeast Hir-Asf1-H3-H4 complex, and 340-370 Å marks the estimated overall width of the human HIRA-nucleosome complex. Human HIRA in complex with Asf1a-H3.3-H4 is estimated to span 340-370 Å. **(d)** Mapping of the HIRA core and CABIN1 density from the human HIRA-Asf1a-H3.3-H4 ab initio model onto the human HIRA-nucleosome structure and the yeast Hir-Asf1-H3-H4 structure. **(e)** Mapping of the yeast Hir core and Hir3 density from the yeast Hir-Asf1-H3-H4 2D class average onto the yeast Hir-Asf1-H3-H4 structure.

**a**

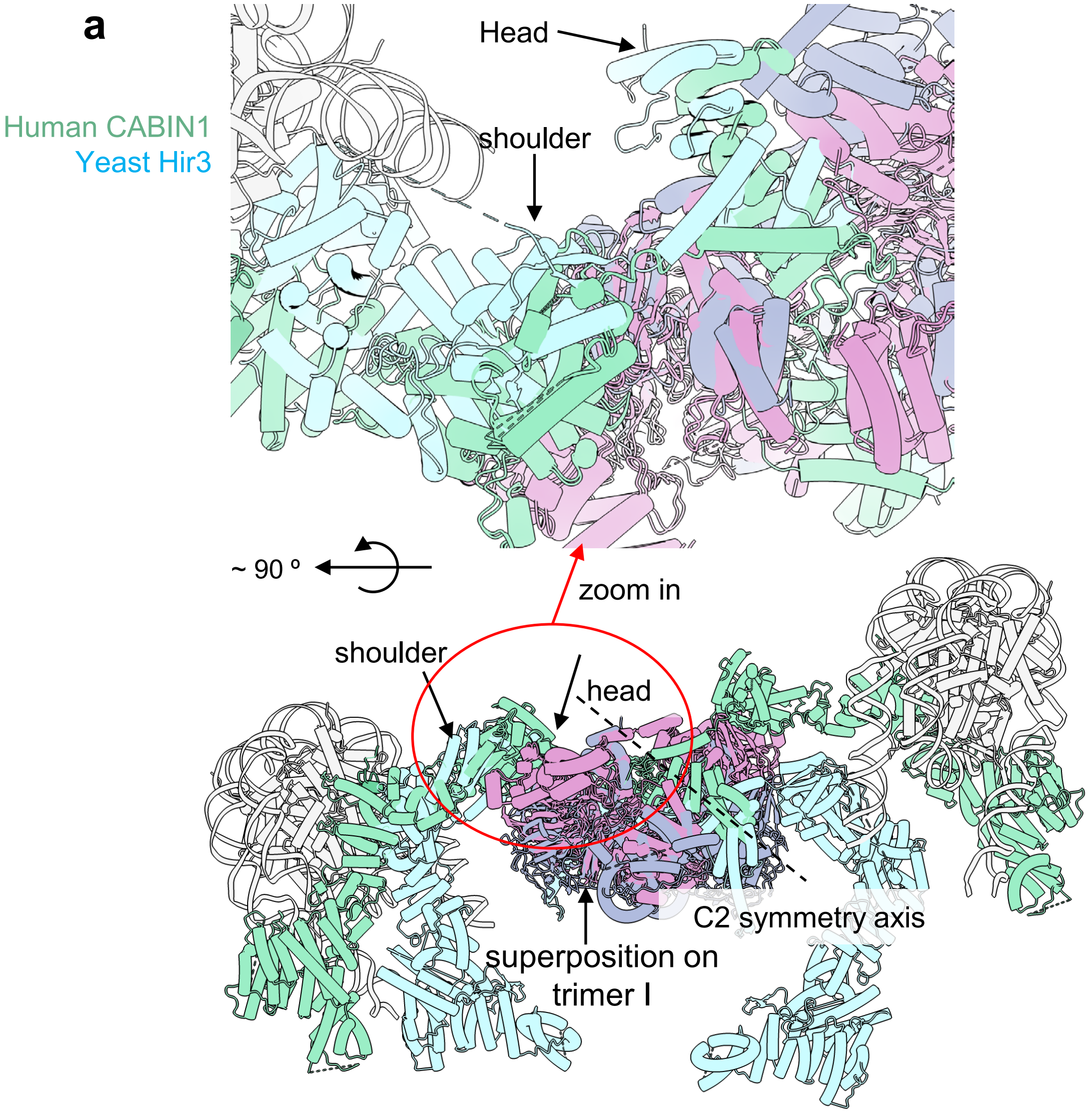


**b**

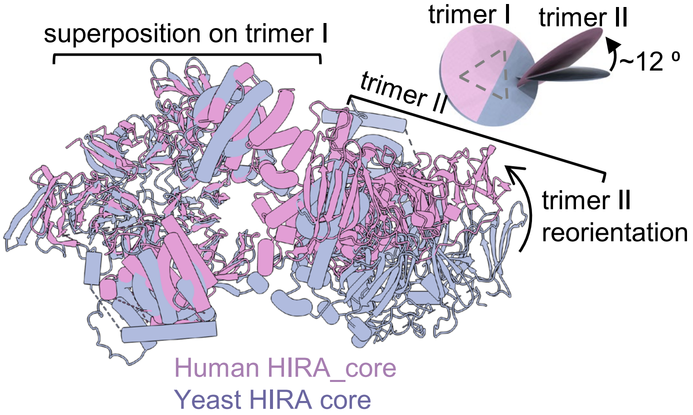


**Supplementary Figure 9. Structural superimposition of the human HIRA-nucleosome complex and yeast Hir-Asf1a-H3-H4, based on HIRA$_s$ trimer I. (a)** Comparison of the location of CABIN1 head and shoulder. Human CABIN1 is shown in green, and the yeast Hir complex is shown in cyan. **(b, c)** Analysis of the relative orientation between the two HIRA trimers in the yeast and human HIRA complex. When the structures are aligned on trimer I, the second HIRA trimer (Trimer II) adopts a different orientation in the human HIRA-nucleosome complex compared to the yeast Hir-Asf1a-H3-H4 structure (PDB 8GHN), corresponding to an approximately 12° reorientation between the two trimers

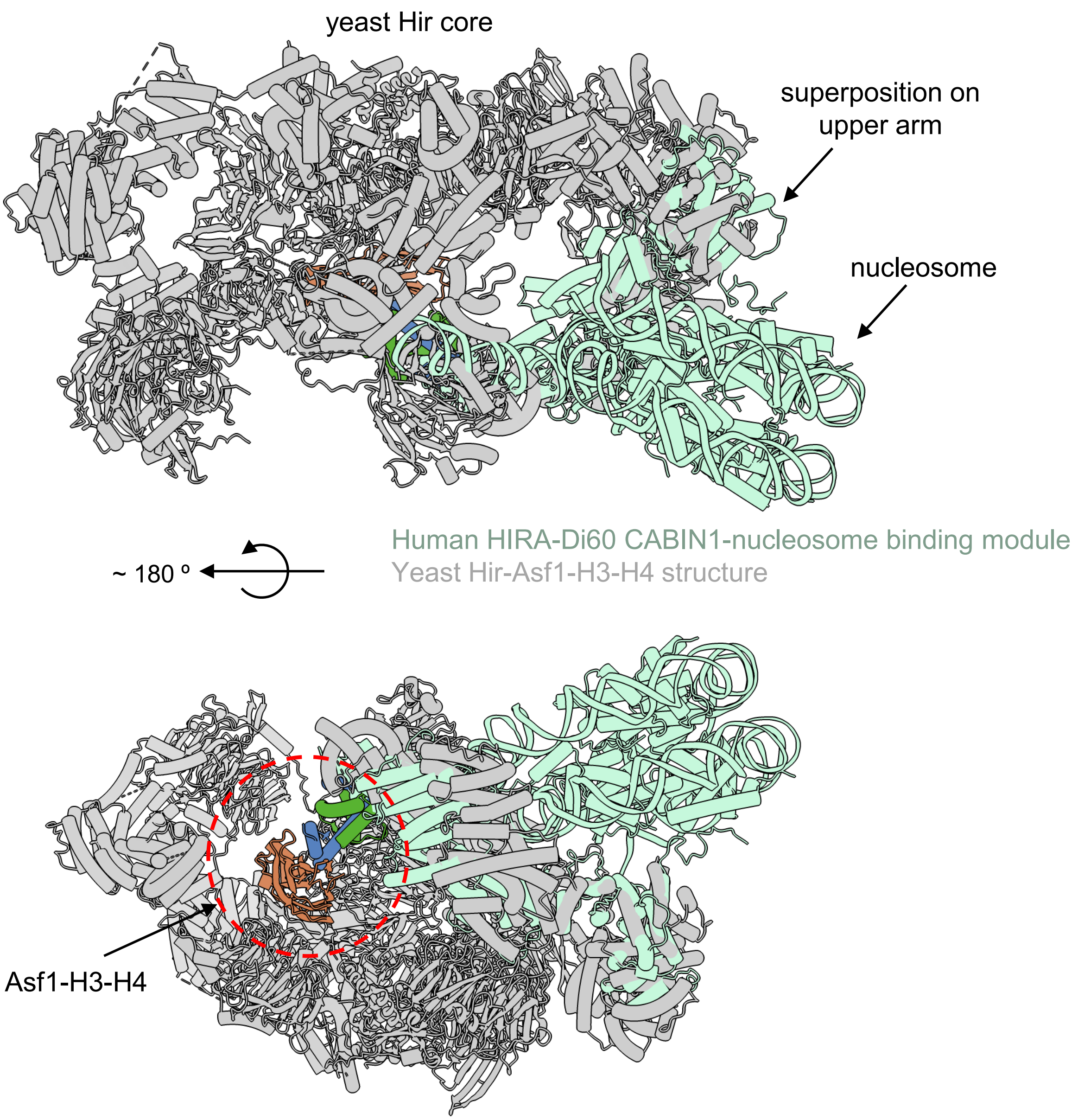


**Supplementary Figure 10. Superposition of HIRA-Di60 CABIN1-nucleosome binding module with the yeast Hir-Asf1-H3-H4 structure.** Alignment is based on the CABIN1 upper arm (CABIN1:936-949 to Hir3:947-960). The CABIN1 nucleosome-binding module from the HIRA-Di60 structure are shown in green.

The yeast structure is shown in grey. The Asf1-H3-H4 complex at the center of the Hir complex is shown in orange, blue and green.

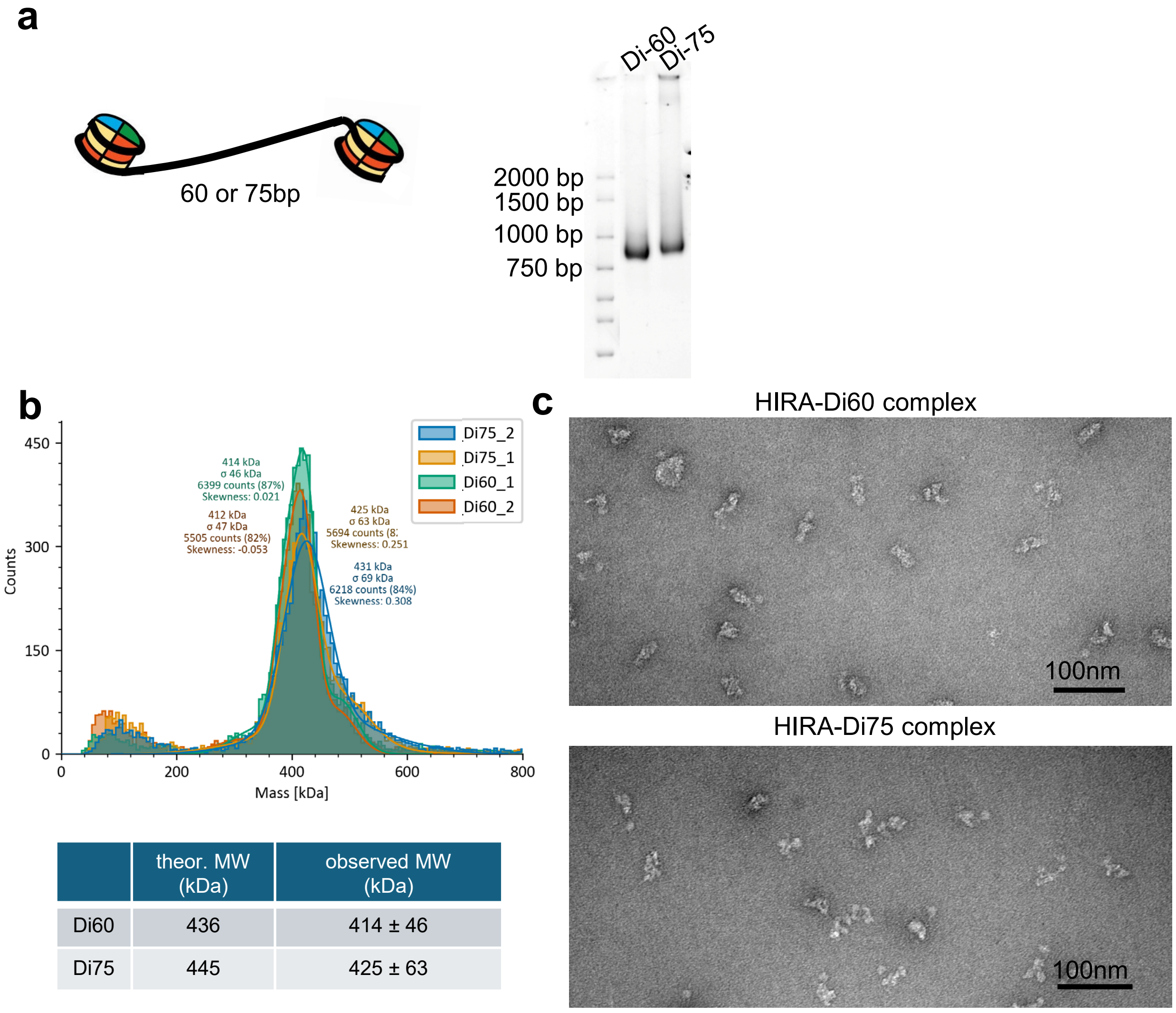


| | theor. MW (kDa) | observed MW (kDa) |
|---|---|---|
| Di60 | 436 | 414 ± 46 |
| Di75 | 445 | 425 ± 63 |

**Supplementary Figure 11: Design and quality assessment of di-nucleosome. (a)** Schematic representation of di-nucleosome substrates containing either a 60-bp or 75-bp linker DNA connecting two 601 nucleosome positioning sequences. Native PAGE analysis confirms successful assembly of the Di60 and Di75 substrates. **(b)** Mass photometry analysis of the assembled di-nucleosome substrates. The observed molecular masses were 414 ± 46 kDa for Di60 and 425 ± 63 kDa for Di75, consistent with the theoretical masses of 436 and 445 kDa, respectively. N=2 **(c)** Representative negative-stain EM micrographs of HIRA bound to Di60 or Di75 substrates. The HIRA-Di60 sample showed a more homogeneous particle population than HIRA-Di75 and was therefore selected for subsequent structural analysis.

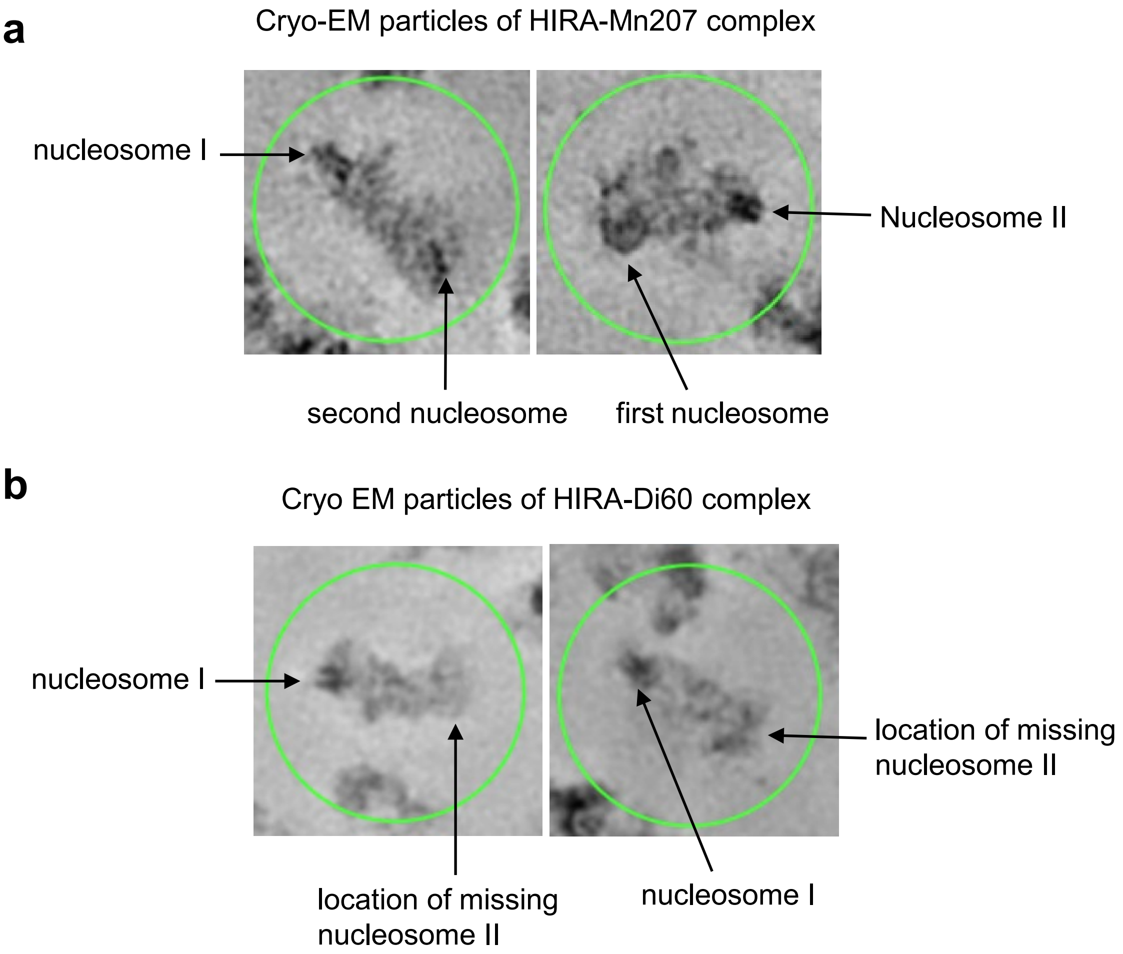


**Supplementary Figure 12: Comparison of cryo-EM particles from HIRA-Mn207 and HIRA-Di60 complex. (**a)HIRA-nucleosome particles; (b) HIRA-Di60 particles. Because DNA has higher contrast under TEM, nucleosomes can be identified based on DNA density. Arrows indicate the positions of the nucleosomes.

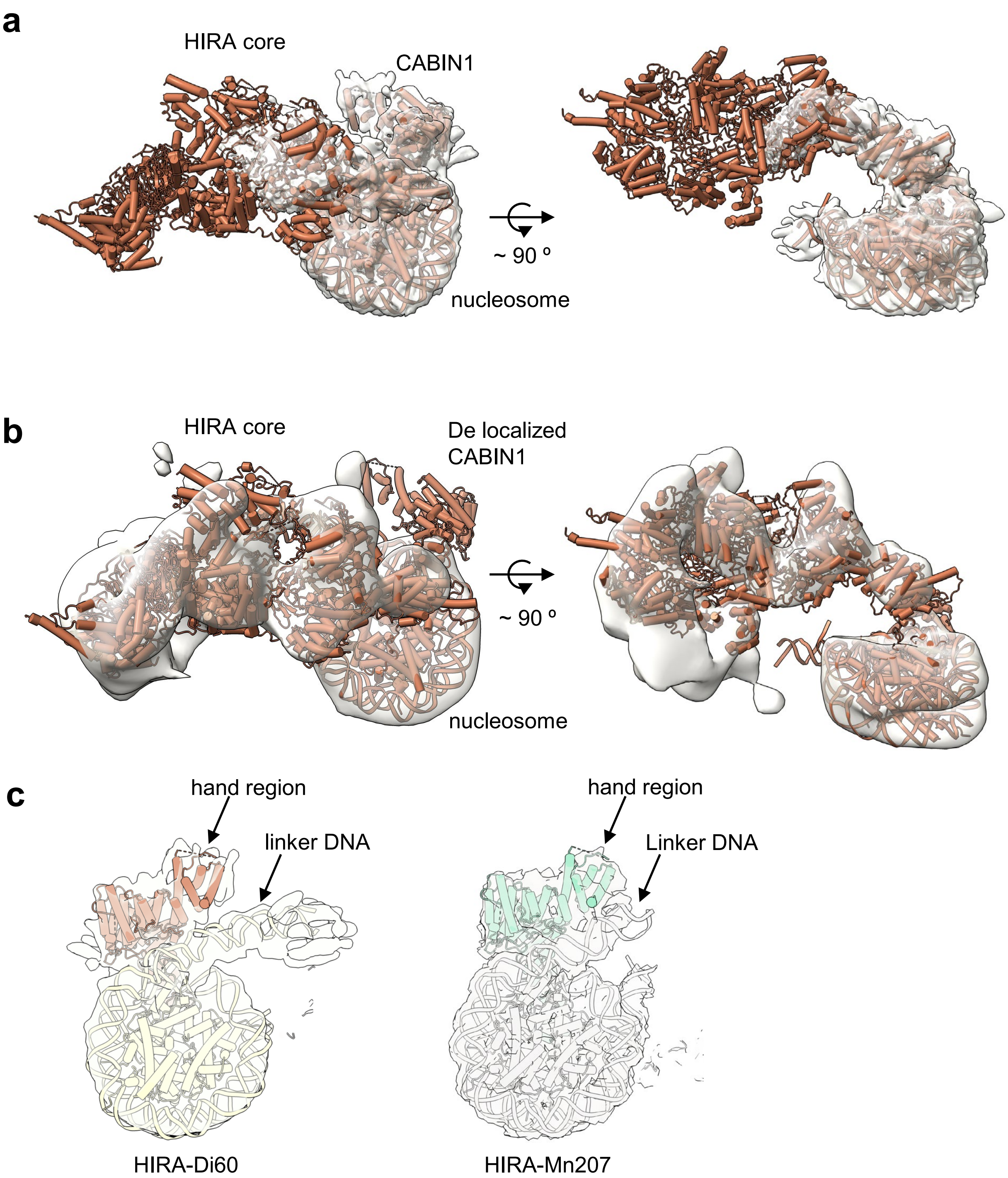


**Supplementary Figure 13: Low-resolution HIRA-Di60 maps supports rigid-body fit of the HIRA core for architectural comparison (a)** Rigid-body fitting of the HIRA-Di60 model into the focused-refinement map of the HIRA-Di60 CABIN1-nucleosome binding module. HIRA-Di60 model only used for HIRA core alignment, not deposited to PDB. HIRA-Di60 CABIN1-nucleosome binding module was deposited to PDB. **(b)** Rigid-body fitting of the HIRA-Di60 model into the overall low-resolution HIRA-Di60 map. **(c)** Cryo-EM density around DNA linker region in both HIRA-Di60 and HIRA-Mn207 structures.

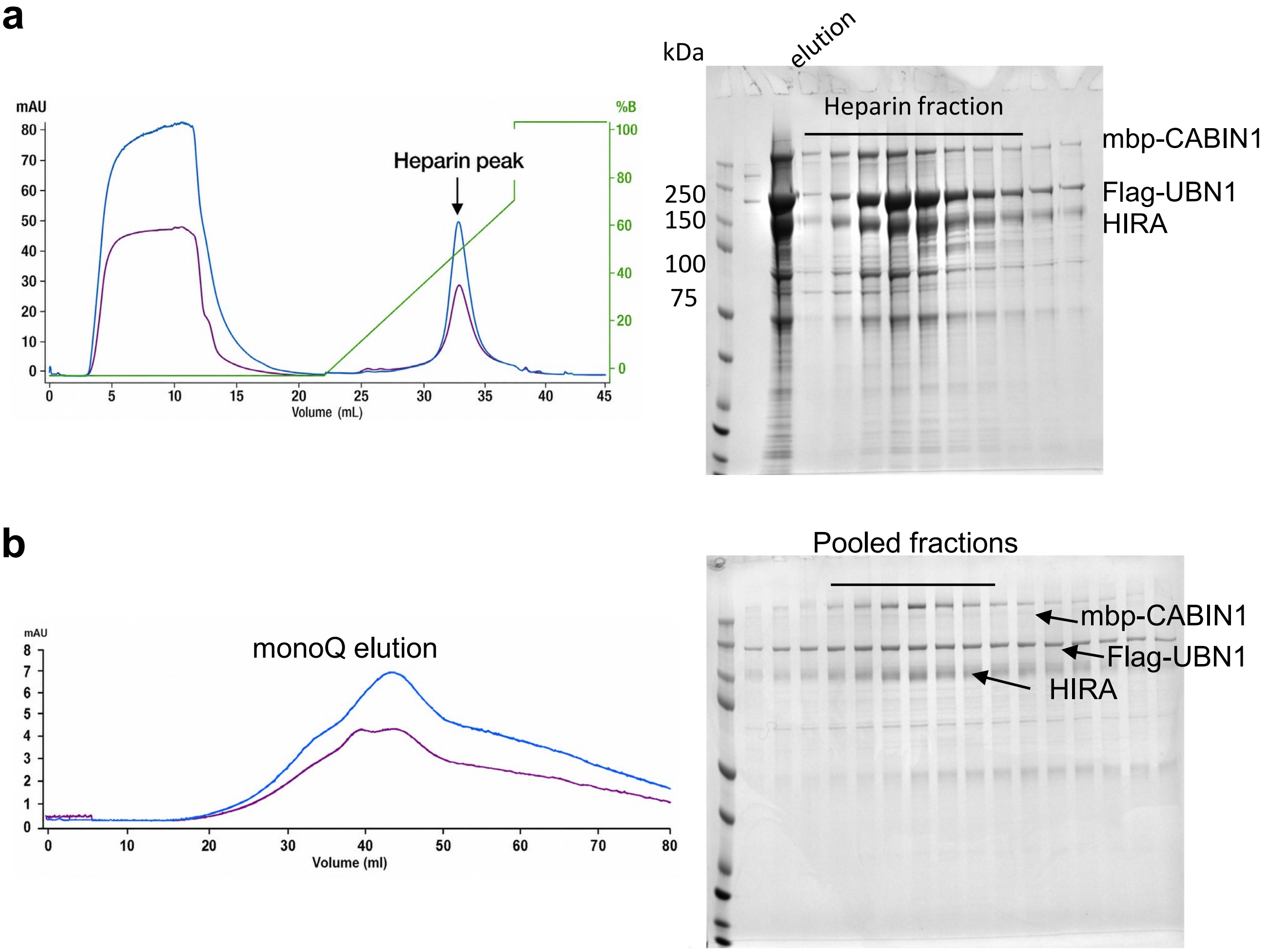


**Supplementary Figure 14: The purification of HIRA complex from mammalian cells.** **(a)** Heparin chromatography profile of the HIRA complex. Fractions across the heparin peak were analyzed by SDS-PAGE (right), and those containing HIRA, FLAG-UBN1, and MBP-CABIN1 were selected for further purification. **(b)** MonoQ chromatography profile of the heparin-purified HIRA complex. Fractions containing all three subunits in similar stoichiometry were identified by SDS-PAGE (right) and pooled for subsequent experiments.

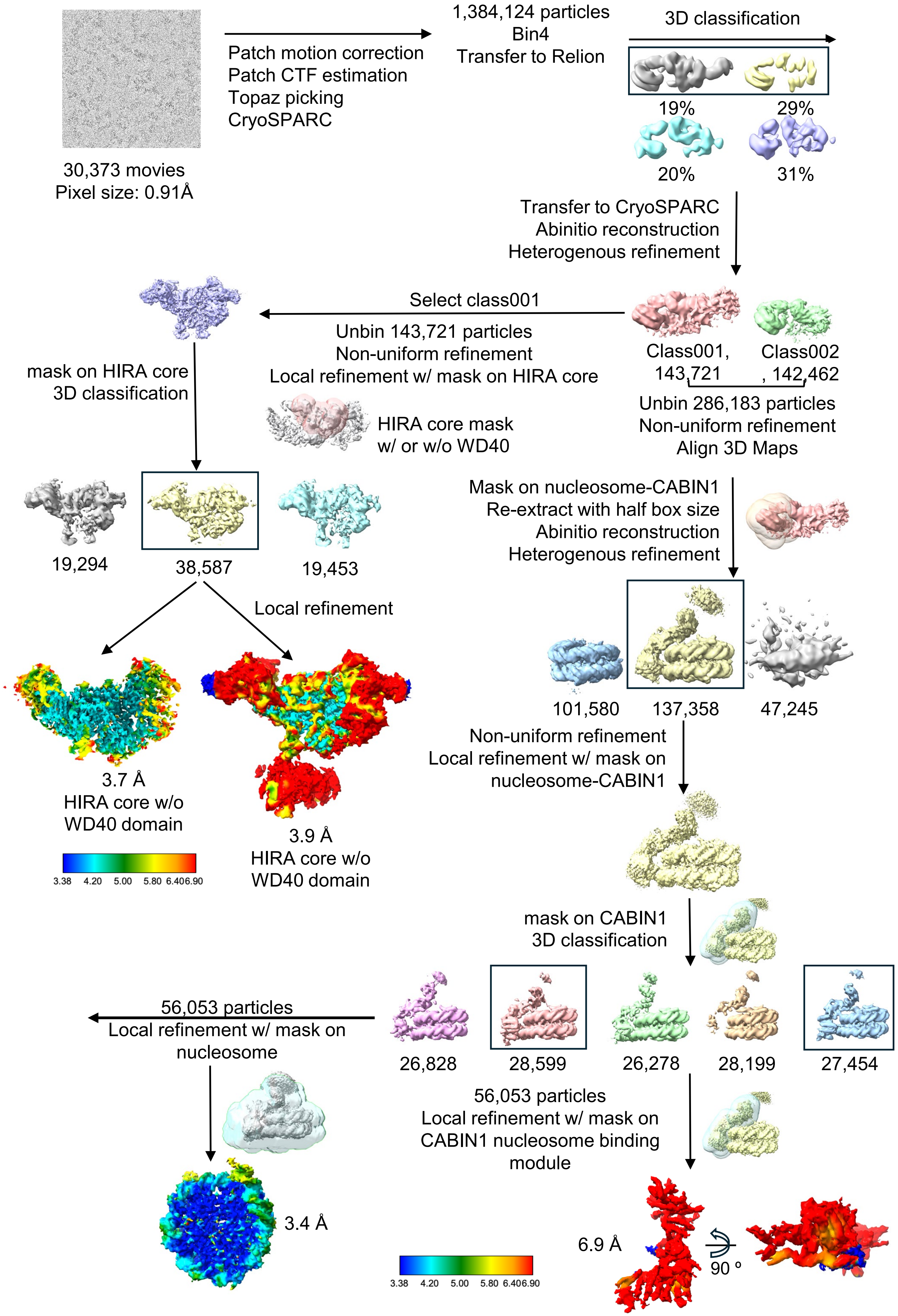


**Supplementary Figure 15:** Data-processing scheme for HIRA-Mn207**.** Flow chart of the Cryo-EM data processing procedure.

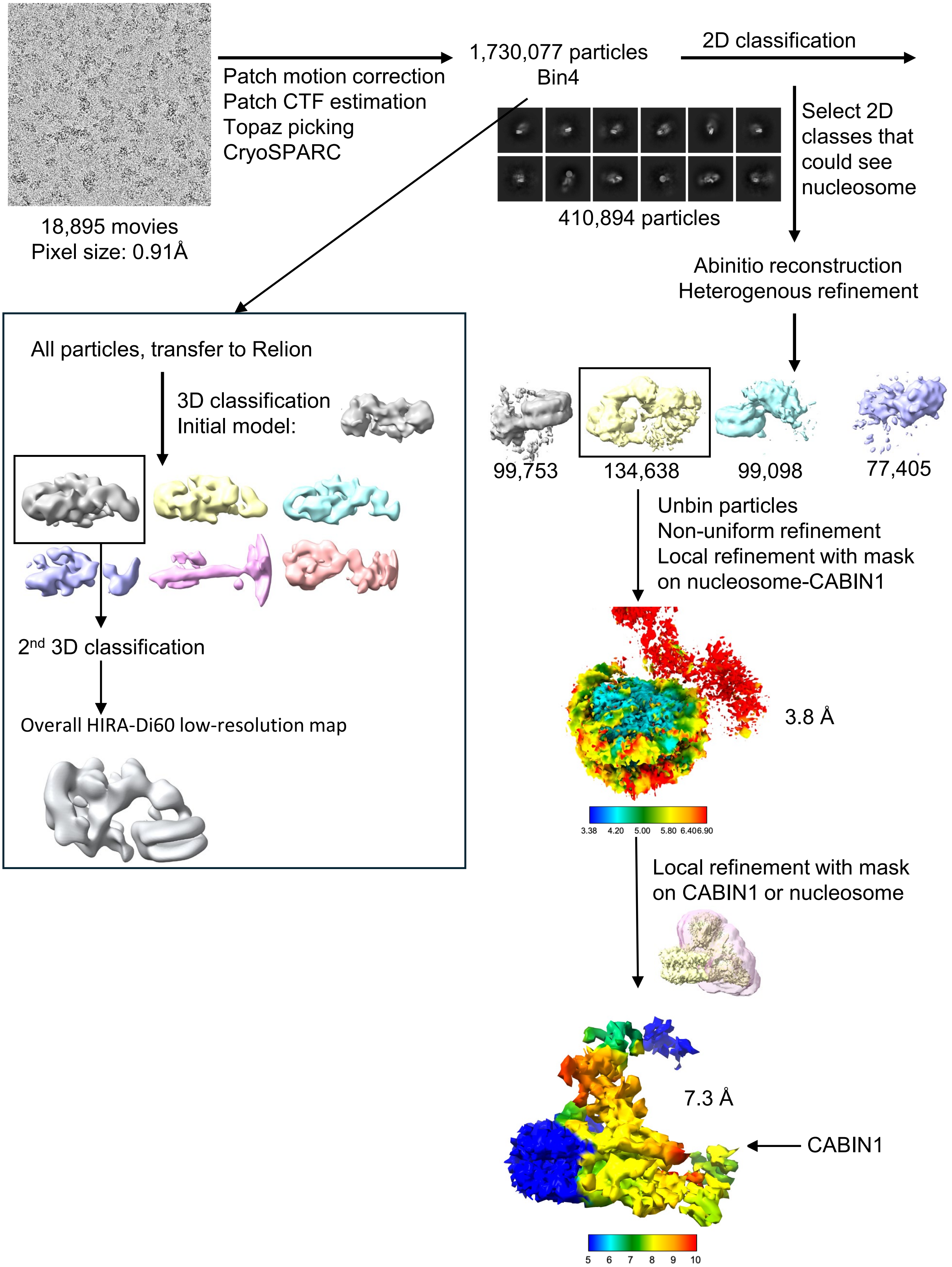


**Supplementary Figure 16: Data-processing scheme of HIRA-Di60 nucleosome complex.** Flow chart of the Cryo-EM data processing procedure.

### HIRA-Mn207 dataset

Locally refine on nucleosome

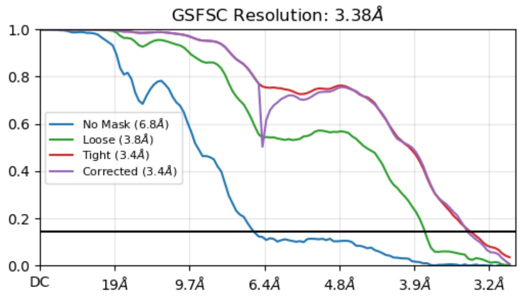


Locally refine on CABIN1

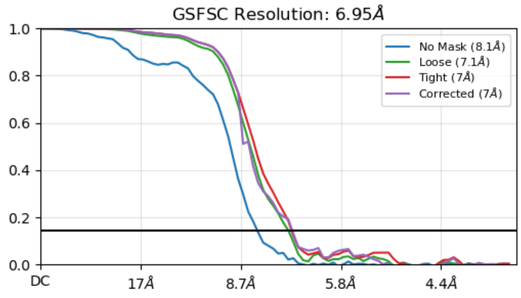


Locally refine on HIRA core w/o WD40 domain

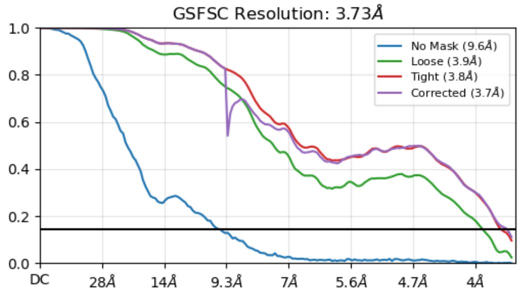


Locally refine on HIRA core w/ WD40 domain

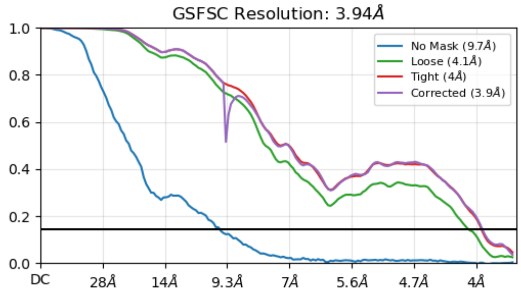


### HIRA-Di60 dataset

Locally refine on nucleosome

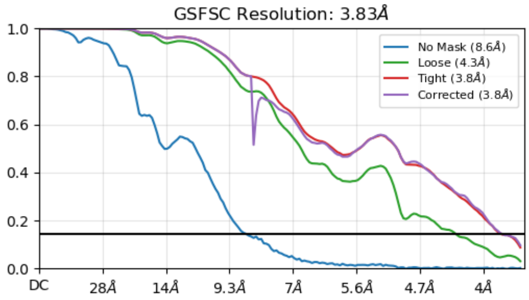


Locally refine on CABIN1

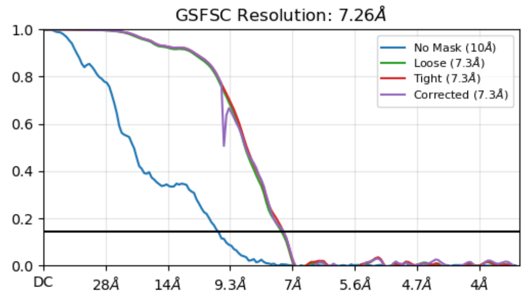


Overall HIRA-Di60 low-resolution map

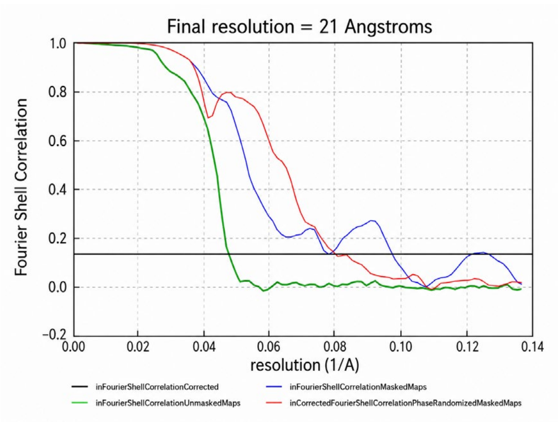


**Supplementary Figure 17: Fourier shell correlation (FSC) curves for all cryo-EM reconstructions**

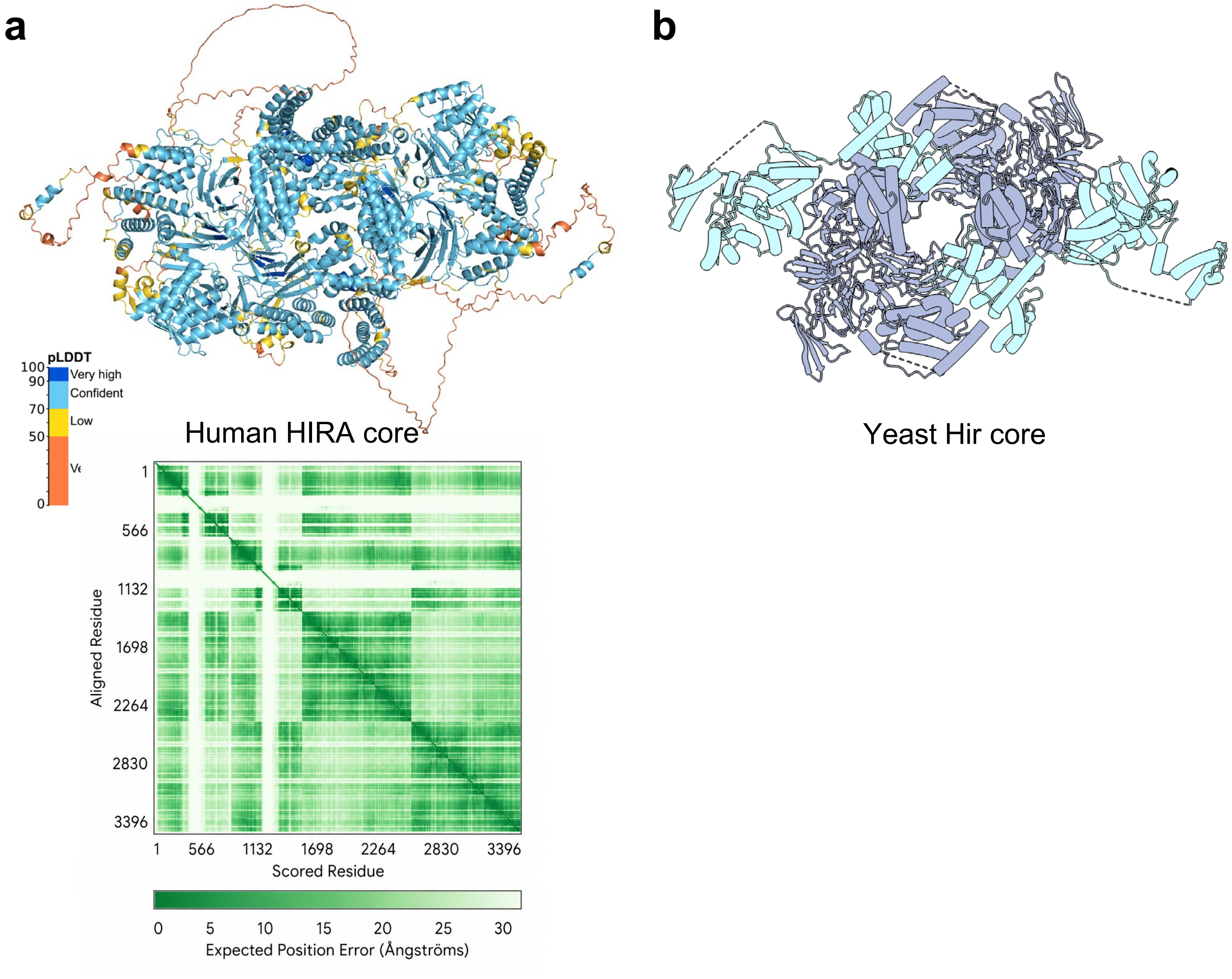


**Supplementary Figure 18 :Alphafold-predicted human HIRA core, with the cryo-EM structure of the yeast Hir core shown as a comparison in the same orientation. (a)** AlphaFold prediction of human HIRAs (680-1017) x 6 and CABIN1 (1-686). pLDDT bar and PAE matrix are shown. **(b)** Cryo EM structure of the yeast Hir core (PDB 8GHN): a complex of Hir1 (418-840) x 2, Hir2 (441-875) x 4 and Hir3 (1-703) x 2 assume a similar shape as the predicted human HIRA core. The two Hir 1-Hir2-Hir2 trimers are colored in purple, the two Hir3 head and shoulder regions are colored in cyan.
