## Supplementary table 1 for "Structure of the human HIRA histone chaperone with a nucleosome suggests a stepwise nucleosome assembly mechanism"

Table S1: Cryo-EM data collection, refinement, and validation statistics of HIRA-Mn207 complex

| Sample | #1 HIRA-Mn207 complex  (EMD-77049: composite map)  (EMD-77044: consensus map)  (EMD-77039: nucleosome-focused refinement map)  (EMD-77041: CABIN1-focused refinement map)  (EMD-77040: HIRA-core-focused refinement map, w/o WD40)  (EMD-77050: HIRA-core-focused refinement map, w/ WD40)  PDB: 13FQ |
| --- | --- |
| Data collection | |
| Microscope | FEI Titan Krios |
| magnification | 130 kx |
| Voltage (kV) | 300 |
| Electron exposure (e-/Å^2^) | 50 |
| Defocus range (μm) | -0.6 ~ -1.4 |
| Pixel size | 0.911 Å |
| Frames per movies | 40 |
| movies | 30,373  EMD-77039: nucleosome = 3.4 Å  EMD-77041: CABIN1= 6.9 Å  EMD-77040: HIRA core w/o WD40 = 3.7 Å  EMDB-77050: HIRA core w/ WD40 = 3.9 Å |
| Map resolution |  |
| Refinement | |
| Bond angles | 0.661 |
| Bond length | 0.004 |
| MolProbity score | 2.65 |
| Clash score | 23.71 |
| Ramachandran plot (%) | |
| Outliers | 0.17 |
| Allowed | 9.97 |
| Favored | 89.86 |
| Rotamer outliers (%) | 0.00 |

Table S2: Cryo-EM data collection, refinement, and validation statistics of HIRA-Di60 complex

| Sample | #2 HIRA-Di60 complex  (EMD-77078: nucleosome-CABIN1 module composite map)  (EMD-77089: HIRA-Di60 overall low-resolution map)  (EMD-77076: nucleosome-focused refinement map)  (EMD-77077: CABIN1-focusd refinement map)  PDB: 13IA |
| --- | --- |
| Data collection | |
| Microscope | FEI Titan Krios |
| magnification | 130 kx |
| Voltage (kV) | 300 |
| Electron exposure (e-/Å^2^) | 50 |
| Defocus range (μm) | -0.6 ~ -1.4 |
| Pixel size | 0.911 Å |
| Frames per movies | 40 |
| movies | 18,895  EMDB-77076: nucleosome-CABIN1 = 3.9 Å  EMDB-77077: CABIN1 = 7.3 Å |
| Map resolution |  |
| Refinement | |
| Bond angles | 0.783 |
| Bond length | 0.006 |
| MolProbity score | 2.18 |
| Clash score | 19.69 |
| Ramachandran plot (%) | |
| Outliers | 0.06 |
| Allowed | 5.75 |
| Favored | 94.19 |
| Rotamer outliers (%) | 0.00 |
